# Error rates in *Q_ST_ –F_ST_* comparisons depend on genetic architecture and estimation procedures

**DOI:** 10.1101/2024.10.28.620737

**Authors:** Junjian J. Liu, Michael D. Edge

## Abstract

Genetic and phenotypic variation among populations is one of the fundamental subjects of evolutionary genetics. One question that arises often in data on natural populations is whether differentiation among populations on a particular trait might be caused in part by natural selection. For the past several decades, researchers have used *Q*_*ST*_ –*F*_*ST*_ approaches to compare the amount of trait differentiation among populations on one or more traits (measured by the statistic *Q*_*ST*_) with differentiation on genome-wide genetic variants (measured by *F*_*ST*_). Theory says that under neutrality, *F*_*ST*_ and *Q*_*ST*_ should be approximately equal in expectation, so *Q*_*ST*_ values much larger than *F*_*ST*_ are consistent with local adaptation driving subpopulations’ trait values apart, and *Q*_*ST*_ values much smaller than *F*_*ST*_ are consistent with stabilizing selection on similar optima. At the same time, investigators have differed in their definitions of genome-wide *F*_*ST*_ (such as “ratio of averages” vs. “average of ratios” versions of *F*_*ST*_) and in their definitions of the variance components in *Q*_*ST*_ . Here, we show that these details matter. Different versions of *F*_*ST*_ and *Q*_*ST*_ have different interpretations in terms of coalescence time, and comparing incompatible statistics can lead to elevated type I error rates, with some choices leading to type I error rates near one when the nominal rate is 5%. We conduct simulations under varying genetic architectures and forms of population structure and show how they affect the distribution of *Q*_*ST*_ . When many loci influence the trait, our simulations support procedures grounded in a coalescent-based framework for neutral phenotytpic differentiation.

## 1 Introduction

Natural selection is a fundamental evolutionary process, shaping genetic variation and the fit of organisms to their environments. Evolutionary biologists have developed a variety of methods for identifying natural selection operating in nature or the laboratory (Vitti et al., 2013; Stern and Nielsen, 2019; Kawecki et al., 2012). In order to understand the action of natural selection, it is crucial to identify cases in which we are confident that selection has occurred.

Going back to the work of Wright (Wright, 1949), evolutionary biologists have often studied natural selection by considering phenotypic differentiation among related populations. If mean levels of a phenotype vary greatly among subpopulations, more than baseline levels of genetic differentiation would lead us to expect, then one explanation is that natural selection has driven the subpopulations to different values of the trait. In the last thirty years, *Q*_*ST*_ –*F*_*ST*_ comparisons have been a major framework for testing hypotheses about natural selection on phenotypes (Whitlock, 1999; Edge and Rosenberg, 2015; Koch, 2019). To perform such a comparison on a single phenotype, one estimates Wright’s fixation index *F*_*ST*_ using data from putatively neutral genetic markers in a set of populations of interest. One then computes an analogous statistic, *Q*_*ST*_ (Spitze, 1993; Prout and Barker, 1993), that measures differentiation on a phenotype, designed to be equal in expectation to *F*_*ST*_ if the phenotype has evolved neutrally. (In fact, the expectation of *Q*_*ST*_ is often slightly less than *F*_*ST*_ (Miller et al., 2008; Edge and Rosenberg, 2015; Koch, 2019).) To rule out environmental explanations for trait differentiation, it is important that *Q*_*ST*_ be estimated from individuals raised in a common garden rather than sampled directly from natural populations (Brommer, 2011; Edelaar et al., 2011; Harpak and Przeworski, 2021; Schraiber and Edge, 2024). *Q*_*ST*_ values much larger than *F*_*ST*_ are consistent with divergent selection driving populations’ phenotypic values apart, perhaps as a result of local adaptation. On the other hand, *Q*_*ST*_ values much smaller than *F*_*ST*_ are consistent with stabilizing selection on a shared optimum or on very similar optima. (We focus here on type I errors in tests of the local adaptation hypothesis.) *Q*_*ST*_ –*F*_*ST*_ comparisons have been widely used to identify selection on phenotypic variation (Whitlock, 2008; Merilä and Crnokrak, 2001; Le Corre and Kremer, 2012).

Notwithstanding their wide use, *Q*_*ST*_ –*F*_*ST*_ comparisons have also faced statistical and conceptual scrutiny (Hendry, 2002; Whitlock, 2008; Edelaar et al., 2011). One issue with *Q*_*ST*_ –*F*_*ST*_ comparisons is ambiguity—there are multiple versions of both *Q*_*ST*_ and *F*_*ST*_, as well as at least two ways of averaging *F*_*ST*_ across loci. Additionally, there are multiple proposed approaches to developing a null distribution for *Q*_*ST*_ . (See Theory and Methods below.) Investigators who use *Q*_*ST*_ –*F*_*ST*_ comparisons implicitly make choices about these dimensions, in addition to choices about experimental design and sampling variation (Whitlock, 2008).

Here, we study how these statistical choices affect the results of *Q*_*ST*_ –*F*_*ST*_ comparisons. We simulate neutral trait variation under a variety of models of population structure and genetic architecture, and we use multiple methods for comparing *F*_*ST*_ and *Q*_*ST*_ . Our results broadly support interpretation of *Q*_*ST*_ –*F*_*ST*_ comparisons in terms of the neutral coalescent, as coalescent-based predictions about which pairings of *Q*_*ST*_ estimator and null distribution will lead to calibrated tests are correct in every case we examine. Encouragingly, the methods that seem to be used most often in the literature are often broadly supported, and our framework explains why these frequent choices often work well.

## 2 Theory and Methods

### 2.1 Theory

When using *Q*_*ST*_ –*F*_*ST*_ comparisons to study trait differentiation, investigators need to make a number of choices. First, one needs to choose a version of *Q*_*ST*_ . Next, one needs to choose a version of *F*_*ST*_, and potentially a way of averaging *F*_*ST*_ values across loci. Finally, one needs to choose a method for generating a null distribution of *Q*_*ST*_ . We discuss each of these decisions in turn, pointing out how the available choices can be interpreted in terms of the coalescent process. For a summary of our notation, see Table 1.

**Table 1.**
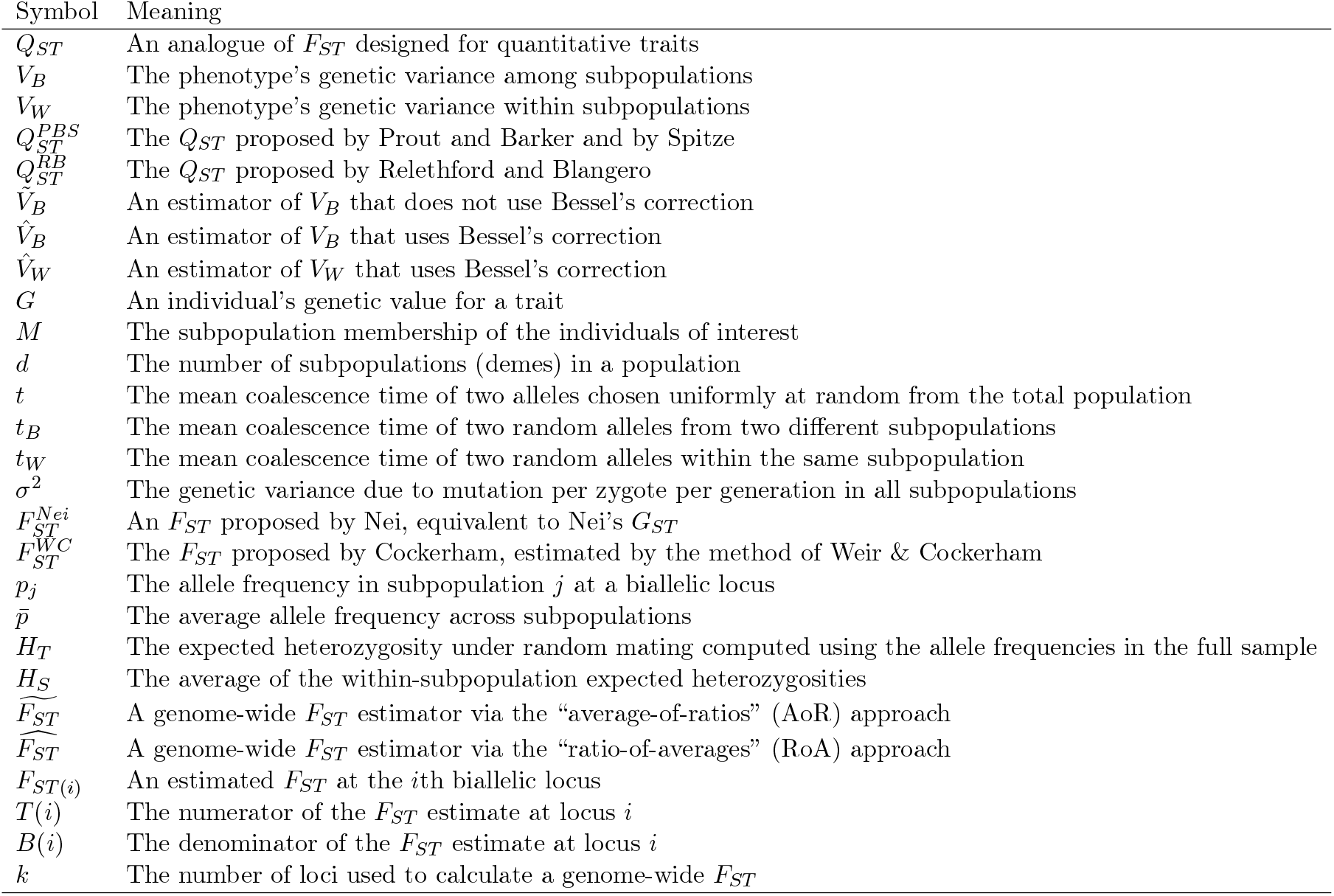
Summary of Notation.

#### 2.1.1 Estimators of *Q*_*ST*_

*Q*_*ST*_ is an analogue of *F*_*ST*_ designed for quantitative traits. For diploids and a single phenotype, it is defined as

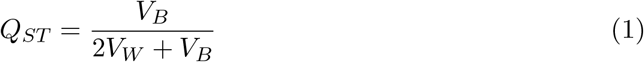

where *V*_*B*_ is the phenotype’s genetic variance among subpopulations and *V*_*W*_ is the genetic variance within subpopulations, that is, the weighted average of the within-subpopulation genetic variances, with weights proportional to the size of each subpopulation. (For general ploidy *ℓ*, the 2 in equation 1 is replaced by *ℓ*. This term is necessary to equilibrate *Q*_*ST*_ with *F*_*ST*_, which can be thought of as a variance proportion for a random draw of a single haploid allele, (Edge and Rosenberg, 2015).)

In general, the genetic variances *V*_*B*_ and *V*_*W*_ are unknown and must be estimated. There are several experimental designs for estimating *V*_*B*_ and *V*_*W*_ involving common gardens. For simplicity, we imagine that individual genetic values for the phenotype are known—or equivalently, that the phenotype is not susceptible to any environmental influence—thus abstracting away from these design considerations. Instead, we focus on two forms of *Q*_*ST*_ estimator proposed independently by three groups in the early 1990s. One estimator was developed independently by Spitze (1993) and by Prout and Barker (1993) and is commonly used in evolutionary biology. The other was proposed by Relethford and Blangero (Relethford and Blangero, 1990; Relethford, 1994) and is more commonly used by evolutionary anthropologists. Following Weaver (2016), we call the version proposed by Prout and Barker and by Spitze 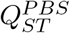, and the version proposed by Relethford and Blangero 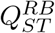 .

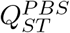 and 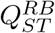 differ according to whether they apply Bessel’s correction to the estimated among-subpopulation genetic variance. That is,

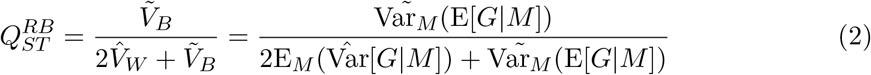

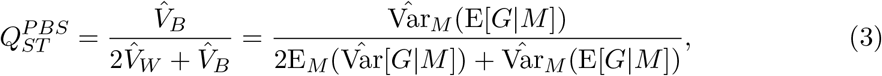

where *G* indicates individual-level genetic value for the trait (i.e. the trait *Y* is conceived as the sum of genetic and environmental components, *Y* = *G* + *E*) and *M* is a variable representing subpopulation membership. Further, 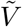 represents an estimator of variance that does not use Bessel’s correction, i.e. for 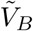, the sum of squared differences between subpopulation means and the grand mean is divided by *d*, the number of demes. In contrast, 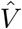 signifies a variance estimator that uses Bessel’s correction. 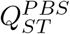 entails Bessel’s correction, dividing the sum of the squared differences between subpopulation means and the grand mean by *d* − 1. Thus, the estimators are very similar when the number of demes *d* is large, but will be quite different for very small numbers of demes. Whitlock (2008) mentions this distinction, writing “It is also essential that the methods used to calculate *F*_*ST*_ and *Q*_*ST*_ both calculate variance among groups in the same way, e.g. by dividing by the number of populations minus one.” But in general it has received little attention, perhaps in part because it is a subtle difference if *d* is large, and in part because 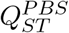 and 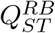 are used by different communities of researchers.

Weaver (2016) showed that 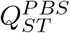 and 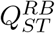 have different interpretations in terms of coalescence times; we follow his exposition in the remainder of this subsection. Let *t* be the mean coalescence time of two alleles chosen uniformly at random from the “total” population, *t*_*B*_ the mean coalescence time of two random alleles from two different subpopulations, and *t*_*W*_ the mean coalescence time of two random alleles within the same subpopulation. Let *σ*^2^ be the genetic variance due to mutation per zygote per generation in all subpopulations. Weaver showed that

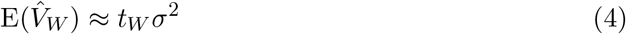

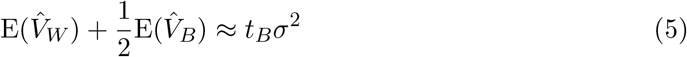

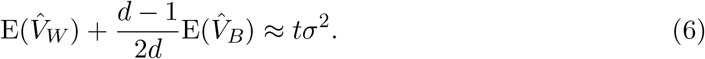

Since 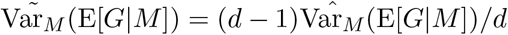, equation 6 can be written as

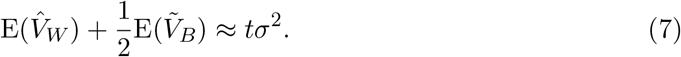

Plugging equations 4 and 7 into the ratio of the expectations of the numerator and denominator of equation 2 gives

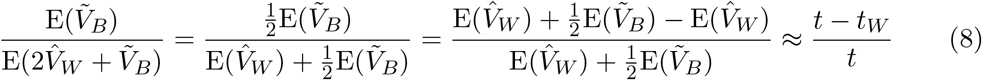

which implies

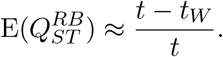

Similarly, combining equations 4–5 with equation 3 gives

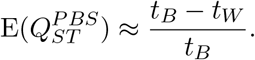

(In both of these equations, the expression on the right is a ratio of the approximate expectations of the numerator and denominator of the 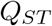 estimator, which is not generally equal to the expectation of 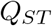, but can be seen as an approximation motivated by a first-order Taylor expansion.)

With large numbers of equally sized demes, *t* ≈ *t*_*B*_, because most random pairs of alleles are from distinct subpopulations. However, with small numbers of demes, it is reasonable to expect that 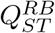 and 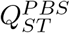 may be most promising when paired with *F*_*ST*_ estimators that estimate the same functions of coalescence times they do under neutrality.

#### 2.1.2 *F*_*ST*_ conceptualizations

Few quantities of interest in evolutionary genetics have inspired more alternative definitions and interpretations than *F*_*ST*_ (Wright, 1949; Nei, 1973; Weir and Cockerham, 1984; Slatkin, 1991; Holsinger and Weir, 2009; Bhatia et al., 2013; Ochoa and Storey, 2021; Goudet and Weir, 2023). *F*_*ST*_ has been variously interpreted as a measure of population differentiation, a “genetic distance” (but see Arbisser and Rosenberg (2020)), an index of the strength of the Wahlund effect on heterozygosity, a correlation of alleles drawn from the same population, an inbreeding coefficient, an estimator of split time or migration rate among populations, an indicator of selection at a locus, a proportion of variance in an indicator variable for allelic type, and a measure of progress toward fixation on different alleles in multiple subpopulations. Here, we do not attempt to encompass the full diversity of approaches to *F*_*ST*_, instead focusing on two versions of *F*_*ST*_ that lead to different interpretations in terms of either variance proportions and coalescence time, and on two methods for averaging *F*_*ST*_ across loci to form a genome-average *F*_*ST*_ .

In this section, we focus on Nei’s *G*_*ST*_ (Nei, 1973), which we call 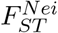, and on Cocker-ham’s (1969; 1973) formulation of *F*_*ST*_, which he called Θ and is estimated by the method of Weir & Cockerham (1984), and which we call 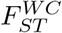. We do not consider descendants of the population-specific *F*_*ST*_ framework developed by Weir & Hill (2002).

Wright defined *F*_*ST*_ in terms of the correlation of a pair of gametes drawn at random from the same subpopulation compared with draws of gametes from the “total” population. The fundamental difference between the approaches of Nei and Cockerham can be understood as stemming from different conceptions of the “total” population. Nei’s definition emerges from an understanding in which the “total” population is the complete sample, that is, the members of all subpopulations sampled. In contrast, Cockerham’s formulation treats the “total” population as an ancestral population from which all the contemporary samples descend. Importantly, in Cockerham’s formulation, we imagine the sampled populations as instances of an evolutionary process of descent from the same ancestor, and *F*_*ST*_ is viewed as a parameter describing that process. This is in contrast to Nei’s formulation, which does not explicitly posit an ancestral population or an evolutionary process, but instead describes the structure of genetic diversity in a sample. This difference is sometimes expressed by saying that the tradition of Nei views *F*_*ST*_ as a statistic, whereas the tradition of Cockerham views *F*_*ST*_ as a parameter (Weir and Cockerham, 1984).

For a set of subpopulations descended from the same ancestral population, Cockerham defined *F*_*ST*_ as a correlation of gametes drawn at random from the same subpopulation compared with pairs of gametes drawn from the population ancestral to the set of subpopulations. Assuming that all subpopulation allele frequencies have drifted independently and by the same amount since their shared ancestor leads to the estimator of Weir & Cockerham (1984). If there are samples of *n* chromosomes from each of *d* subpopulations, then the Weir & Cockerham estimator for the *i*th biallelic locus simplifies to

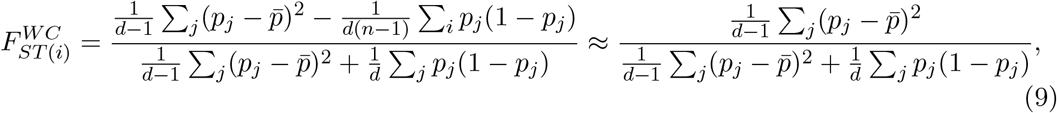

where *p*_*j*_ is the allele frequency in subpopulation 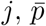 is the average allele frequency across subpopulations, and the approximation holds if the sample size per subpopulation (i.e. *n*) is large.

In contrast, Nei’s *F*_*ST*_ analogue, which he labeled *G*_*ST*_, is defined as

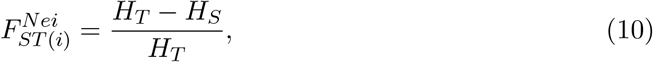

where *H*_*T*_ is Nei’s “gene diversity” (i.e. the expected heterozygosity under random mating) computed using the allele frequencies in the full sample, and *H*_*S*_ is the average gene diversity within subpopulations. Thus, at the *i*th biallelic locus, and with equal sample sizes per subpopulation, Nei’s *F*_*ST*_ can be estimated as

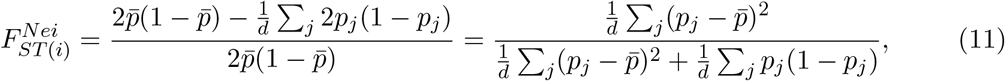

where the second equality comes from the fact that 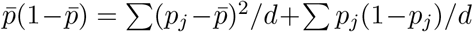 (Ehm, 1991). Potentially adding to the confusion over *F*_*ST*_, Nei (1986) suggested a second form of *F*_*ST*_, which he labeled 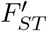, in which the numerator of equation 11 is multiplied by *d/*(*d* − 1), rendering the numerator equal to that of the right side of equation 9. Bhatia and colleagues (2013) refer to this alternative 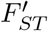 as Nei’s *F*_*ST*_, whereas our references to Nei’s *F*_*ST*_ are to his original formulation from 1973, and we do not consider 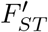 further.

Comparing equations 9 and 11 reveals that Nei’s 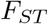 estimator would be approximately equal to Weir & Cockerham’s estimator (assuming large and equal sample sizes per subpopulation) if the terms corresponding to among-subpopulation variation (i.e. the numerator and the first term of the denominator) were divided by *d* − 1 instead of *d*. Thus, they will be approximately equal for large numbers of subpopulations. This view also reveals a correspondence between these two forms of *F*_*ST*_ and the forms of *Q*_*ST*_ considered above. Specifically, both Weir & Cockerham’s 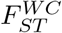 and the Prout–Barker–Spitze 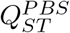 apply Bessel’s correction to the estimator of variance among groups (as noted in passing by Whitlock (2008)), whereas Nei’s 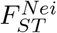 and Relethford & Blangero’s 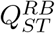 do not apply Bessel’s correction.

The correspondence between 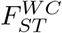 and 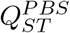, on one hand, and 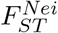 and 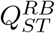 is also apparent when considering their interpretation in terms of average coalescent times. As pointed out by Slatkin (1991), for low mutation rates, Nei’s 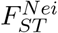, expressed in terms of probabilities of identity, has a low-mutation-rate limit of (*t* − *t*_*W*_)*/t*, where *t* is the average pairwise coalescence time for gametes drawn uniformly from the population at large, and *t*_*W*_ is the average coalescence time for pairs of gametes drawn from the same subpopulation. This expression in terms of coalescence times exactly matches that for 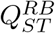 above. Similarly, Slatkin (1993) pointed out that the analogous limit for Weir & Cockerham’s 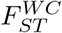 is (*t*_*B*_ − *t*_*W*_)*/t*_*B*_, where *t*_*B*_ is the average coalescence times for pairs of gametes drawn from different subpopulations. This expression matches that for 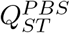, a correspondence pointed out by Weaver (2016).

Thus, theoretical considerations, whether viewed from the perspective of variance partitioning or coalescence times, lead us to expect that Relethford and Blangero’s 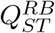 is comparable with Nei’s 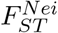 and that the Prout–Barker–Spitze 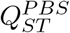 is comparable with Weir & Cockerham’s 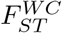. Because the most general motivations for comparison of *Q*_*ST*_ and *F*_*ST*_ are based on coalescent arguments (Whitlock, 1999; Koch, 2019), the coalescent argument takes special importance. Because both sets of estimators become more similar for large numbers of subpopulations, we might also predict that the differences matter most for small *d*.

#### 2.1.3 Averaging *F*_*ST*_ estimators

Given a choice of a single-site estimator of *F*_*ST*_, there are two major strategies for estimating genome-wide *F*_*ST*_ . Perhaps the most obvious approach is simply to take the average of the *F*_*ST*_ values at each locus. Because *F*_*ST*_ is a ratio, this is sometimes called the “average-of-ratios” (AoR) approach, and can be written as

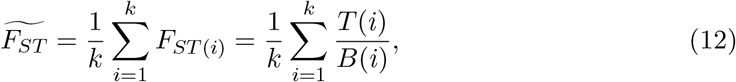

where *T* (*i*) is the numerator and *B*(*i*) is the denominator of the *F*_*ST*_ estimate at locus *i*, and *k* is the number of loci. The other major approach is to sum separately the numerators and denominators of the *F*_*ST*_ estimates at all loci and then report their ratio as the final estimate. This is sometimes called a “ratio-of-averages” (RoA) approach and can be written as

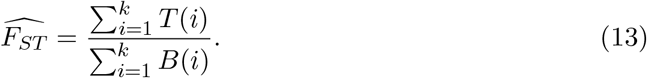

Whereas the average-of-ratios estimator is an unweighted average of the single-locus *F*_*ST*_ estimates, the ratio-of-averages estimator is a weighted average, where the weights are the denominators of the single-locus *F*_*ST*_ estimates, which themselves are generally estimates of the total variation at the locus. That is, the ratio-of-averages estimator can be written as

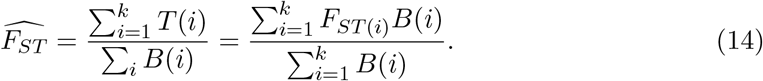

Empirically, when loci with low minor allele frequency are included in estimates of *F*_*ST*_, the average-of-ratios estimator tends to produce smaller estimates than the ratio-of-averages estimator (Bhatia et al., 2013). This observation makes sense—ratio-of-averages *F*_*ST*_ estimators down-weight loci with low minor allele frequencies, since they also have low total heterozygosity, and *F*_*ST*_ at loci with low minor allele frequencies is mathematically constrained to be small (Jakobsson et al., 2013; Alcala and Rosenberg, 2017).

As ratio estimators, both the ratio-of-averages and average-of-ratios approach may produce biased estimates, since the expectation of a ratio is not generally equal to the ratio of the expectations of its numerator and denominator. Weir & Cockerham (1984) recommended a ratio-of-averages approach to averaging *F*_*ST*_ . More recently, Guerra & Nielsen (2022) studied sequence-based estimators of *F*_*ST*_ . Their results imply that, with two subpopulations, the average-of-ratios approach will typically be biased downward as an estimator of *F*_*ST*_, interpreted as a function of coalescence times. Using a downwardly biased genome-wide *F*_*ST*_ estimator could result in an excess of *Q*_*ST*_ tests that produce spurious evidence of local phenotypic adaptation.

#### 2.1.4 Proposed null distributions for *Q*_*ST*_

The reason that the estimator of *F*_*ST*_ matters for *Q*_*ST*_ − *F*_*ST*_ comparisons is that we wish to form a null distribution that describes the behavior of *Q*_*ST*_ under neutrality. We consider three broad approaches that have been proposed in the literature. First, we consider the Lewontin–Krakauer distribution, a re-scaled *χ*^2^ distribution parameterized to have an expectation equal to a genome-wide estimate of *F*_*ST*_ (Lewontin and Krakauer, 1973). We consider versions of the Lewontin–Krakauer distribution with expectations equal to *F*_*ST*_ estimates coming from either the Nei or Weir–Cockerham estimators, and from genome-wide averages of *F*_*ST*_ based on either the ratio-of-averages or average-of-ratios approach. The Lewontin–Krakauer distribution was derived under the assumption of a star-like population tree. This suggests that it may work poorly for demographic models with spatial structure or other departures from starlike demography, although it has also been suggested to be fairly robust to such deviations in some contexts (Beaumont, 2005).

The Lewontin–Krakauer distribution was developed as an approximation to the distribution of single-locus *F*_*ST*_ values. Thus, an alternative approach is to use the realized distribution of single-locus *F*_*ST*_ values as a null distribution for *Q*_*ST*_ . This approach is well-justified for single-locus traits and has been shown to perform well with simulated traits governed by a small number of loci (Whitlock, 2008). We consider the distribution of single-locus *F*_*ST*_ for all loci or for common variants only (see below).

Finally, we tested an approach recently recommended by Koch (2019). Koch’s method involves identifying the covariance matrix expected among subpopulations evolving neutrally for the genetic component of a quantitative trait, then simulating multivariate normal random variables with that covariance matrix and computing *Q*_*ST*_ values from them to form a null distribution of *Q*_*ST*_ . Given any pair of subpopulations, their covariance is computed on the basis of mean pairwise coalescent times under neutrality within and between the subpopulations. (See equation 10 in Koch 2019. As we discuss below, Koch’s expressions are consistent with the Relethford & Blangero version of *Q*_*ST*_ .)

### 2.2 Simulation methods

We sought to simulate neutral genetic variation with many subpopulations under a variety of demographic models. Diffusion-based approaches to compute the approximate joint site-frequency spectrum (SFS) (Gutenkunst et al., 2009; Jouganous et al., 2017) are limited to fewer demes than we require. We thus used a coalescent approach to generate approximate joint site-frequency spectra (Nielsen, 2000; Excoffier et al., 2013). With large numbers of demes, the joint SFS is high dimensional and has too many entries to estimate the frequency of rare allele-frequency configurations accurately by simulation. Nonetheless, the approach allows us to draw genetic variants with allele frequencies that are consistent with the demographic models we study. A schematic description of our protocol is shown in Figure 1A.

**Figure 1.**
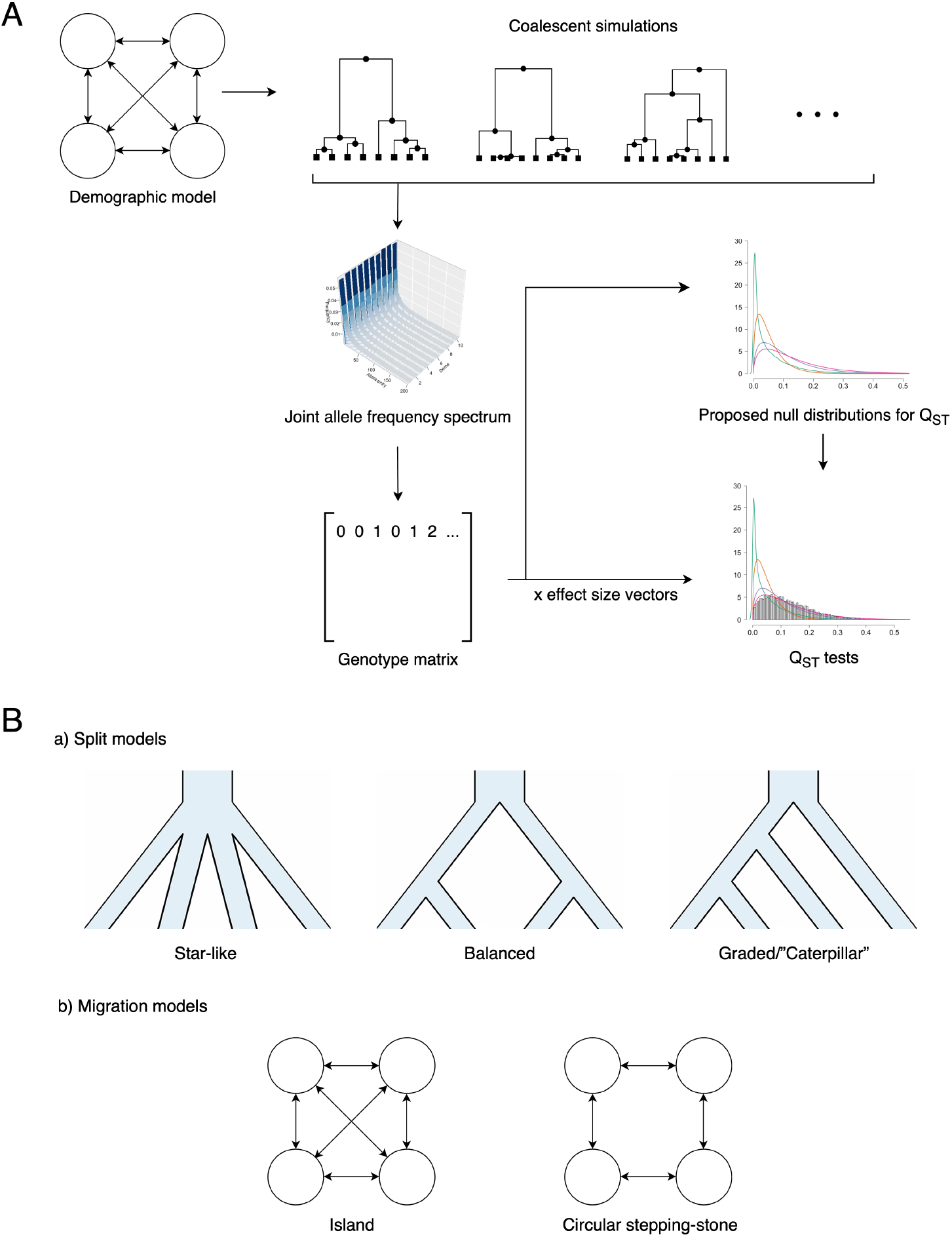
Schematic figure of (A) simulations and (B) demographic models. We simulated independent coalescent trees and used the branch lengths to compute an approximate joint site-frequency spectrum for each demographic model. Demographic modes included three scenarios involving splits among subpopulations (star-like, balanced, and graded/caterpillar) and two scenarios involving migration among subpopulations (island and circular stepping-stone).

#### 2.2.1 Joint site-frequency spectrum approximations

We ran simulations to generate independent coalescent trees obeying each of the demographic models we studied and approximated the joint allele frequency spectrum on the basis of tree branch lengths. This procedure has been used previously (Nielsen, 2000; Excoffier et al., 2013). More formally, we ran *R* simulations and estimated the joint site frequency spectrum entry corresponding to the existence of *s* = (*s*_1_, *s*_2_, …, *s*_*d*_) copies of an allele in demes 1, 2, …, *d* as:

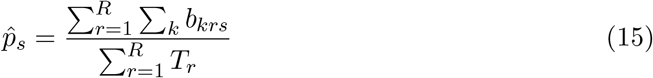

where *b*_*krs*_ represents the length of the *k*_*th*_ branch in the *r*_*th*_ simulated tree that is compatible with joint SFS entry *s*. That is, *b*_*krs*_ is the length of a branch that has exactly *s*_1_ descendants in subpopulation 1, *s*_2_ descendants in subpopulation 2, and so on. *T*_*r*_ is the total branch length of the *r*th simulated tree.

We used msprime (Baumdicker et al., 2022) to simulate 5,000 independent coalescent trees for each demographic setting studied. The branch lengths of every tree were processed by a custom script to allow subsequent computation of equation 15. We did not apply mutations to the simulated trees, instead simulating mutations later via sampling from the estimated joint SFS.

#### 2.2.2 Demographic models

Broadly, we examined two types of demographic models (Figure 1B)—those in which differentiation among subpopulations occurs because subpopulations split from each other in the recent past and do not subsequently exchange migrants (“split models”) and those in which differentiation among long-separated subpopulations reaches an equilibrium value because of constant exchange of migrants (“migration models”).

We examined three kinds of topologies for split models: star-like, in which all subpopulations split from an ancestor at the same time in the past; balanced, i.e. a symmetric, bifurcating tree; and graded/caterpillar, a bifurcating tree in which every split produces one subpopulation that does not split again (except the most recent split, which produces two such subpopulations). In all split models, we set the effective population size to be the same in every branch of the population tree. Among these, the star-like topology is of note because it reflects the assumptions used in the derivation of the Lewontin–Krakauer distribution, as well as those invoked in deriving the Weir–Cockerham estimator of *F*_*ST*_ .

Among migration models, we examined an island model, in which migrants from a given island are equally likely to migrate to any other island, and a circular stepping-stone model, in which migrants from a given island can only migrate to one of its two immediate neighbors. The circular stepping-stone model induces spatial structure that departs strongly from the star-like assumptions used to derive the Lewontin–Krakauer distribution (Koch, 2019).

We simulated each demographic scenario with 2, 4, 8, and 16 subpopulations with 100 diploid individuals sampled per subpopulation respectively. Effective population size *N*_*e*_ per deme was set to 1000 and demographic parameters (split time or migration rates) were adjusted to achieve a values of (*t* − *t*_*W*_)*/t* (which should approximate the expected value of 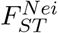) of 0.02, 0.1, or 0.25 across unlinked loci. Theoretical *F*_*ST*_ calculations for each model and scenario are provided in supplementary text.

#### 2.2.3 *Q*_*ST*_ − *F*_*ST*_ comparisons

We compared the distribution of *Q*_*ST*_ to several proposed null distributions. We simulated genotypes first—these genotypes served both to produce single-locus *F*_*ST*_ estimates and, once assigned random effect sizes, to produce individual values of the genetic component of a quantitative trait. For each demographic history, we simulated 20000 random loci according to the approximate joint site-frequency spectrum. A genotype matrix was then produced by randomly pairing these alleles within subpopulations to form sampled individuals. We calculated 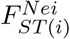and 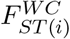 for each locus. Then, we calculated ratio-of-averages and average-of-ratios estimates of genome-wide 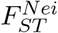 and ratio-of-averages estimates for the effect-size standard deviation is inversely proportional to genome-wide 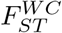 to use as input for parameterizing the Lewontin–Krakauer distribution.

We compared the proposed null distributions with *Q*_*ST*_ distributions of simulated phenotypes. We first generated effect size vectors with entries drawn from various distribution families. An effect size vector is a vector indicating a random subset of loci assigned with randomly drawn effect sizes. Effect sizes were drawn from Gaussian, Uniform, and Laplace distributions with expectation 0 and variance 1. We also tested effect sizes drawn from an “alpha model” with *α* = −1 (an allele-frequency-dependent Gaussian distribution in which the effect-size standard deviation is inversely proportional to 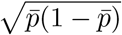, where 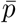 is the mean allele frequency across the total population). We note that the alpha model is not a neutral model, and with a single population, *α* = −1 emerges when there is very strong stabilizing selection on a single trait (Schraiber et al., 2024). Nonetheless, we simulated under the assumption that effect sizes are assigned with respect to average allele frequency, but without respect to differences in frequency among subpopulations given the average frequency.

We simulated traits with 1, 10, 100, or 1000 loci with non-zero effect sizes. Individual phenotypic values were generated by taking the dot product of the effect-size vector with a vector of individual genotypes. We calculated 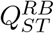 and 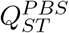 according to equation 2 and 3 for each of 10,000 simulated traits. We measured type I error rates for comparisons against every proposed null distribution of *Q*_*ST*_ . A nominal threshold of *α* = 0.05 used for assessing Type I error rate across all demographic scenarios.

## 3 Results

### 3.1 Ratio-of-averages *F*_*ST*_ approximates the theoretically expected functions of coalescence time

We simulated independent coalescent trees and used the ratio of branch lengths on the tree collection to approximate three joint allele frequency spectra per demographic model, with the value of (*t*−*t*_*W*_)*/t* (which corresponds to 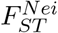) set to 0.02, 0.1, or 0.25. Figure S1 shows that across all models, ratio-of-average estimators of 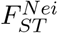 applied to all loci accurately estimated (*t* − *t*_*W*_)*/t*. (Similarly, ratio-of-averages 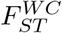 estimated (*t*_*B*_ − *t*_*W*_)*/t*_*B*_ accurately, and were therefore larger on average than (*t* − *t*_*W*_)*/t*, as expected.) In contrast, average-of-ratios estimators always gave smaller values on average. These results change somewhat when loci are selected either on the basis of being common in one target subpopulation (Figure S2) or on average across the total population (Figure S3).

### 3.2 Mean *Q*_*ST*_ appears bounded from above by *F*_*ST*_ under neutrality if the chosen *F*_*ST*_ and *Q*_*ST*_ correspond in terms of coalescence times

We investigated the behavior of 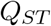 estimates under various demographic scenarios. For each phenotype, we calculated 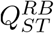 and 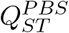 with effect sizes drawn from several distribution families, i.e. normal, uniform, and Laplace distributions. In these simulations across various types of effect sizes, *Q*_*ST*_ estimates show similar patterns (Figure S4). Figure 2 shows results when effect sizes are sampled from a normal distribution. Grey lines show (*t* − *t*_*W*_)*/t*, the function of coalescence times corresponding to 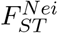. As expected, mean values of 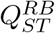 are bounded from above by (*t* − *t*_*W*_)*/t*, though for traits influenced by small numbers of loci, they are substantially lower than this upper bound, as observed previously (Edge and Rosenberg, 2015). Mean values of 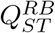 were also smaller than (*t* − *t*_*W*_)*/t* for small numbers of demes.

**Figure 2.**
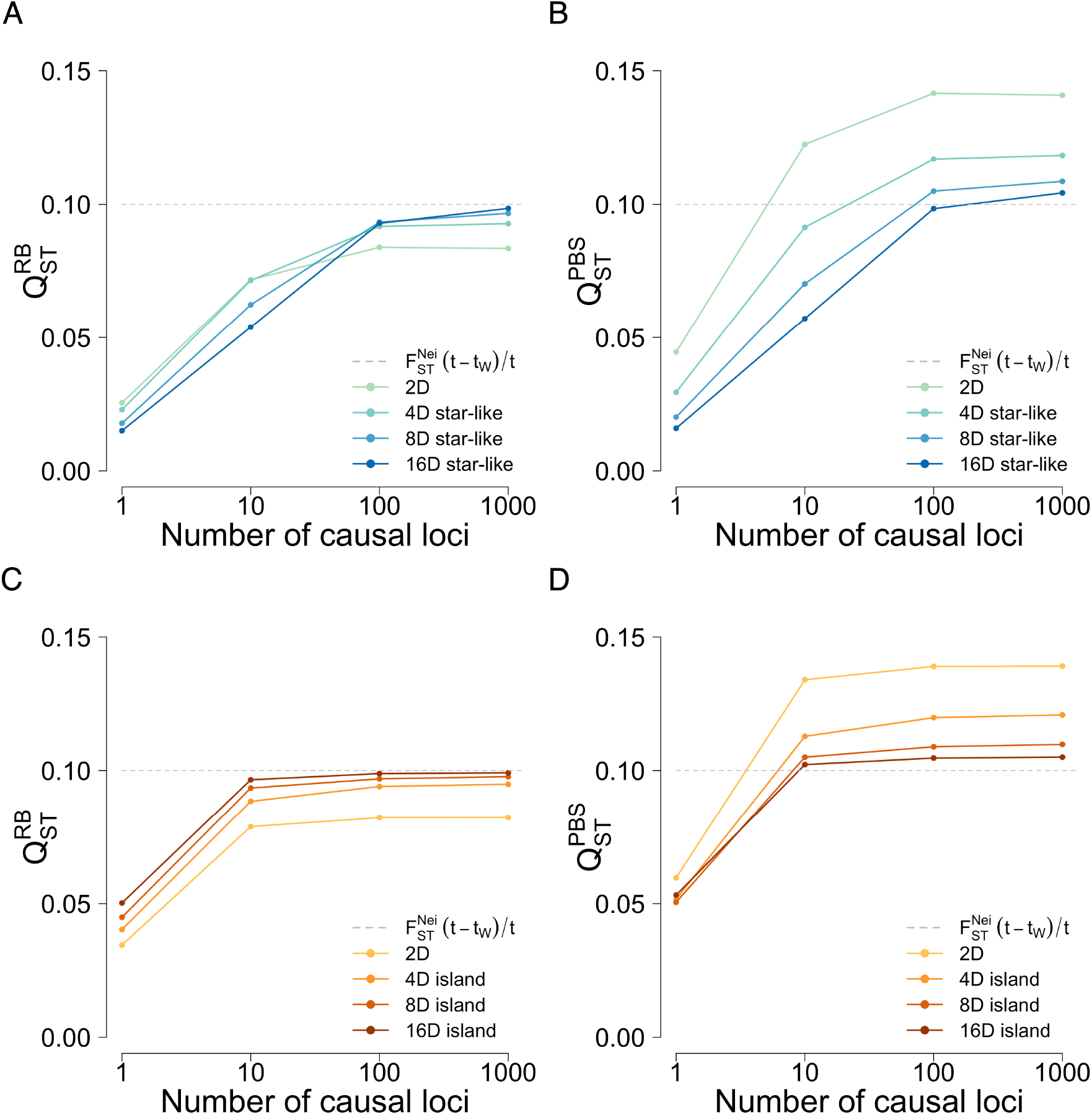
The behavior of mean *Q*_*ST*_ estimates in selected demographic models. Effect sizes were randomly sampled from a Gaussian distribution with variance 1 to generate phenotypic values. Mean *Q*_*ST*_ estimates were calculated across 1000 simulated traits with (*t* − *t*_*W*_)*/t* (i.e. the function of coalescent times estimated by 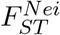) equal to 0.1. The curves in each panel show the behavior of (A) 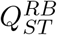 in 2D, 4D, and 8D star-like split models, (B) 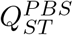 in 2D, 4D, and 8D star-like split models, (C) 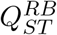 in 2D, 4D, and 8D island models, and (D) 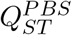 in 2D, 4D, and 8D island models.

Unlike 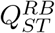, mean values of 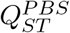 were somewhat larger than (*t* − *t*_*W*_)*/t*, particularly for small numbers of demes. This is again expected, as 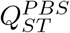 applies Bessel’s correction to the among-population variance in the numerator, causing it to be substantially larger than 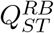 for small numbers of demes. As shown in supplementary Figure S5, the mean value of 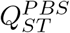 is not larger than (*t*_*B*_ − *t*_*W*_)*/t*_*B*_, the function of coalescence times to which 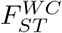 corresponds.

### 3.3 Single-locus *F*_*ST*_ distributions match *Q*_*ST*_ distributions for monogenic traits

We next examined the distribution of *Q*_*ST*_ compared with the distribution of single-locus *F*_*ST*_, considering all variable loci irrespective of allele frequency. Figure 3 shows the distribution of single-locus 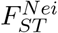 values compared with 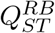 values for simulated traits influenced by 1, 10, 100, or 1000 unlinked loci under a star-like, eight-deme split model. Unsurprisingly, when the simulated phenotype is influenced by one genetic locus, the distributions match closely—in this case, the *Q*_*ST*_ values are equivalent to single-locus *F*_*ST*_ values. However, when the number of loci influencing the trait is larger, the distributions no longer match. Importantly, in these simulations, all loci are equally likely to contribute to the trait, meaning that most single-locus traits will be controlled by relatively low-frequency loci, and so will not vary much either between or within subpopulations. This scenario is perhaps not reflective of most empirical studies, in which traits are likely to be chosen for study in part because they display substantial genetic variance. Figures S6, S7, S8, and S9 show similar results comparing 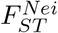 and 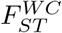 with 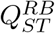 and 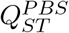.

**Figure 3.**
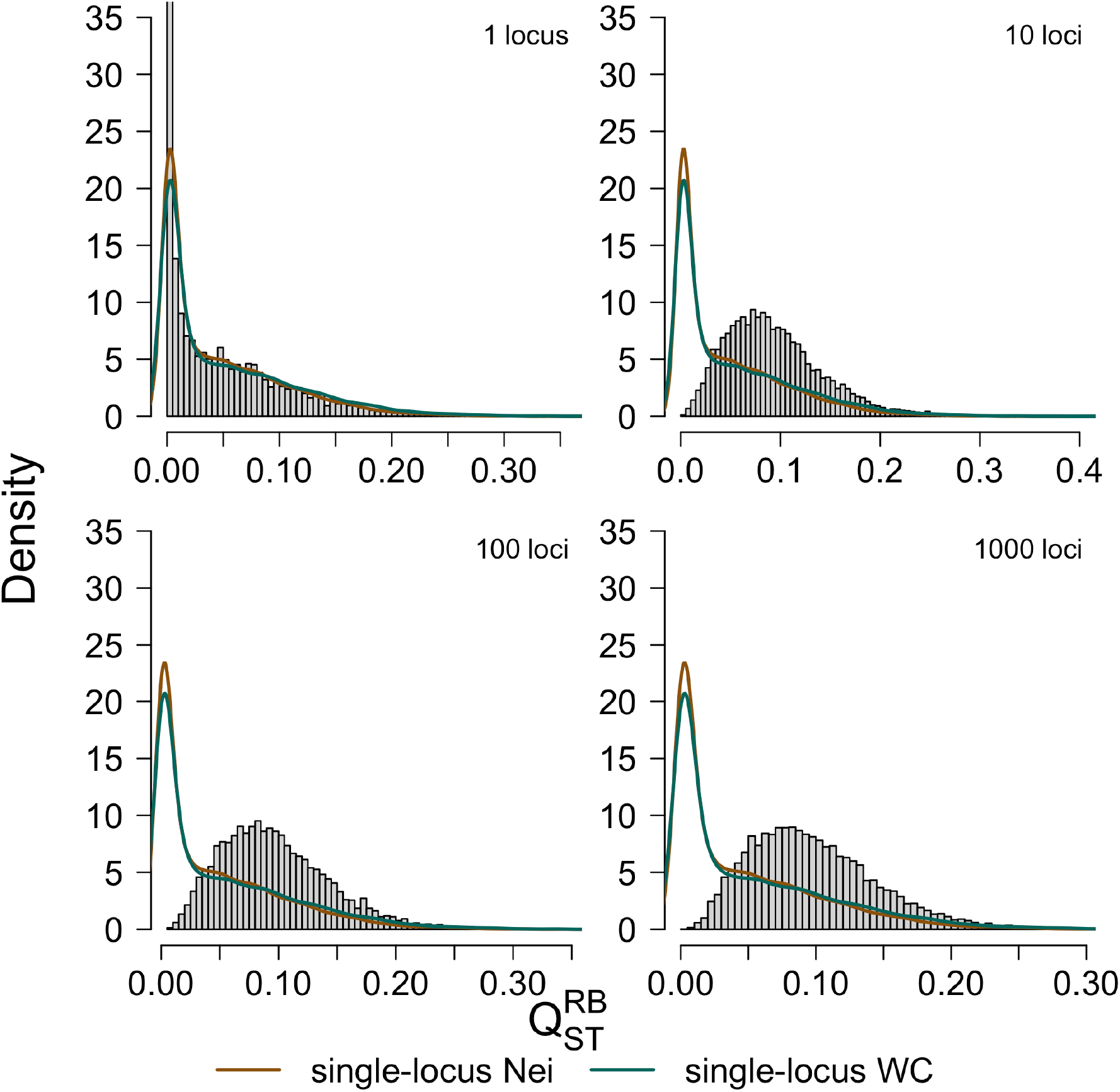
Single-locus *F*_*ST*_ density curves vs. *Q*_*ST*_ distributions across genetic architectures: eight-deme island models. We compared two null distributions (the single-locus 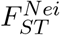 and 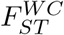 density curves, using all variable loci) with neutral 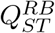 distributions. Each *Q*_*ST*_ distribution included 10,000 traits with 1, 10, 100, or 1000 causal loci. The panels show the results for an eight-deme island model. Effect sizes were randomly sampled from a Gaussian distribution with variance 1. The value of (*t* − *t*_*W*_)*/t* was 0.1.

### 3.4 The Lewontin–Krakauer null works well for polygenic traits without spatial structure if the coalescence interpretation matches

Next, we considered the Lewontin–Krakauer distribution as a null distribution for *Q*_*ST*_ . The Lewontin–Krakauer distribution is a scaled *χ*^2^(*d* − 1) distribution, where the scaling ensures that the expectation of the Lewontin–Krakauer distribution is equal to a genome-wide *F*_*ST*_ . Thus, the performance of the Lewontin–Krakauer distribution depends on the type of genome-wide *F*_*ST*_ estimator used to parameterize it.

Figure 4 shows the fit to *Q*_*ST*_ values from simulated traits of the Lewontin–Krakauer distribution parameterized by either ratio-of-averages or average-of-ratios *F*_*ST*_ values. Parameterizing the Lewontin–Krakauer distribution with average-of-ratios estimators of global *F*_*ST*_ always leads to a poor fit to the distribution of *Q*_*ST*_ . Because average-of-ratios estimators are biased downward as estimators of (*t* − *t*_*W*_)*/t* or (*t*_*B*_ − *t*_*W*_)*/t*_*B*_, they lead to Lewontin–Krakauer distributions centered on low values of *Q*_*ST*_, and these null distributions therefore lead to many false positives (Figures S10, S11, S12, S13, and Table S2).

**Figure 4.**
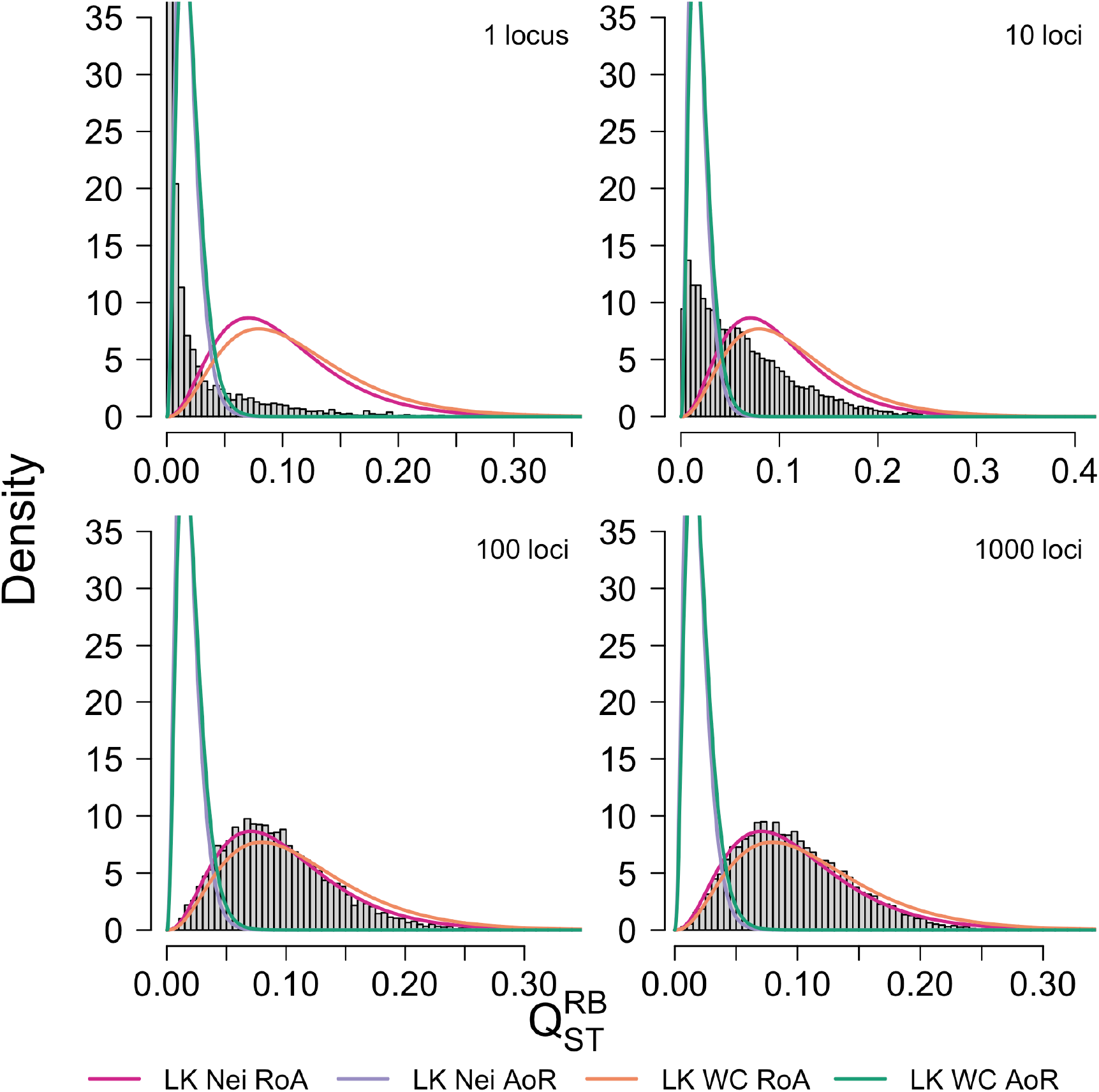
Lewontin-Krakauer null vs. *Q*_*ST*_ distributions across genetic architectures: eight-deme star-like split models. We compared the Lewontin–Krakauer distribution parameterized by either ratio-of-average or average-of ratios estimates of genome-wide 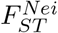 or 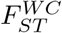 to neutral distributions of 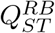. Each *Q*_*ST*_ distribution included 10,000 traits with 1, 10, 100, or 1000 causal loci. The panels show results for an eight-deme star-like split model. Effect sizes were randomly sampled from a Gaussian distribution with variance 1; the value of (*t* − *t*_*W*_)*/t* was 0.1.

However, for polygenic traits, the Lewontin–Krakauer distribution often fits the distribution of neutral *Q*_*ST*_ values well, provided that it is parameterized by a ratio-of-averages

*F*_*ST*_ estimate that matches the definition of *Q*_*ST*_ used. Specifically, the Lewontin–Krakauer distribution fits the neutral distribution of 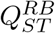 when it is parameterized by a ratio-of-averages estimator of 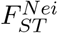, and it matches 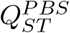 when it is parameterized by a ratio-of-averages estimator of 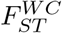, under both the migration and split models (Figures S10, S11, S12, and S13). Both of these choices produce calibrated or slightly conservative tests for local adaptation. However, if 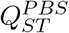 is parameterized by 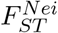, the test is anti-conservative, and if 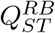 is parameterized by 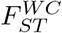, the test is unnecessarily conservative (Table S2). These differences become very small as the number of demes increases.

### 3.5 Lewontin–Krakauer null fails for spatially structured populations with many demes

The original argument for the Lewontin–Krakauer distribution as an approximate distribution for single-locus *F*_*ST*_ assumed a star-like population tree (Lewontin and Krakauer, 1973). Recently, Koch (2019) noticed that the Lewontin–Krakauer distribution is a poor null distribution for *Q*_*ST*_ values from populations with strong spatial structure. The results shown in Figure 5 agree with those of Koch. In circular stepping-stone models with few demes, the Lewontin–Krakauer distribution is an acceptable approximation to the distribution of *Q*_*ST*_ under neutrality, producing conservative *p* values with four demes and only slightly anti-conservative *p* values with eight demes. However, when there are 16 demes, the Lewontin–Krakauer distribution is too symmetric and too strongly peaked at its mode, leading to type I error rates of approximately 10% when the nominal rate is 5% for polygenic traits.

**Figure 5.**
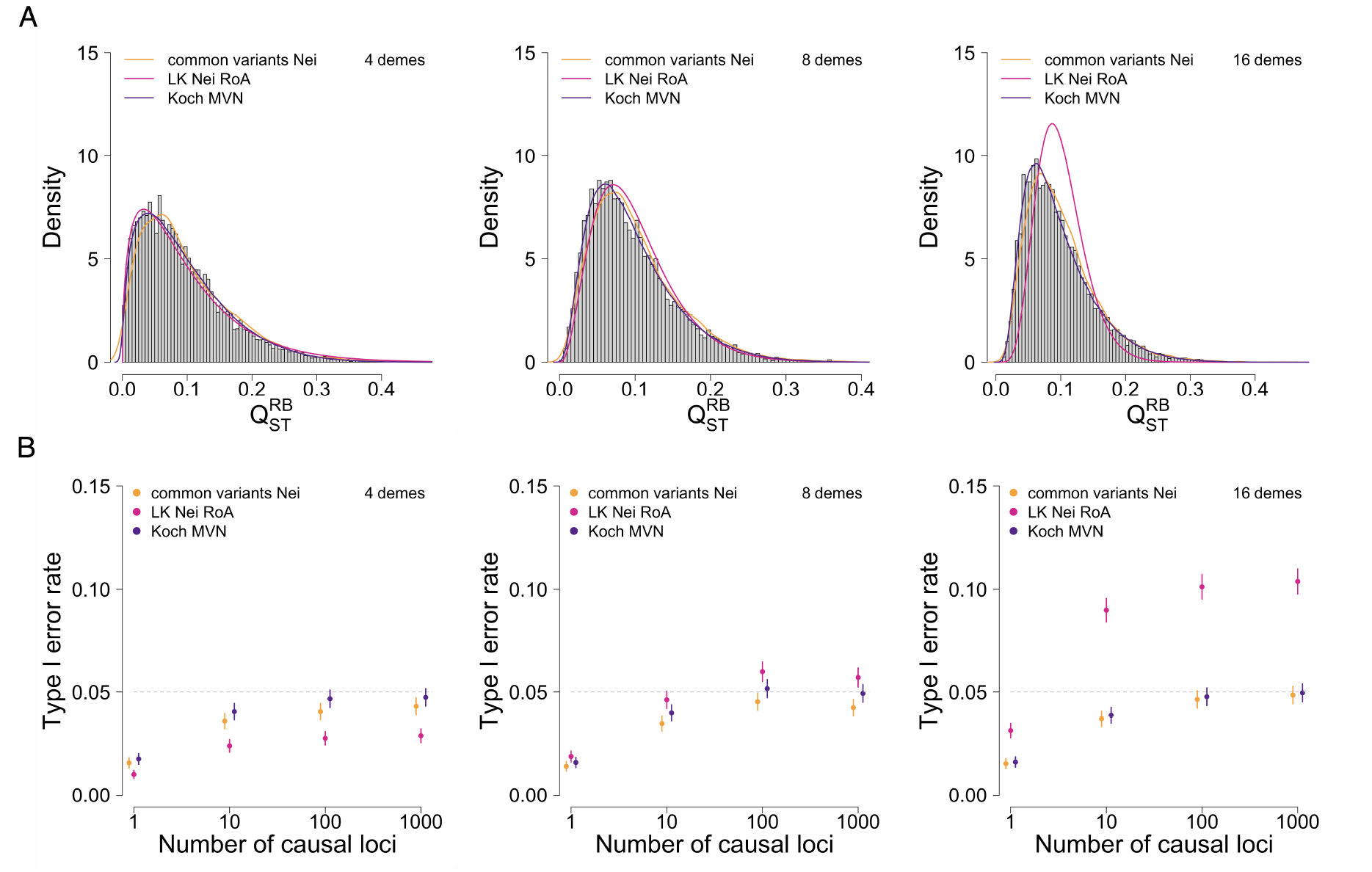
Multiple nulls vs. *Q*_*ST*_ distributions across genetic architectures: four-deme, eight-deme, and sixteen-deme circular stepping-stone models. We compared three different null distributions—from the Lewontin–Krakauer distribution, from single-locus *F*_*ST*_ values from common variants, and from Koch’s (2019) multivariate normal procedure—with neutral 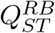 values simulated under circular stepping-stone models. Each *Q*_*ST*_ distribution included 10,000 traits with 1000 causal loci. The panels show the results of (A) three proposed nulls compared to *Q*_*ST*_ distributions and (B) type I error rates in *Q*_*ST*_ –*F*_*ST*_ comparisons of four-deme, eight-deme, and sixteen-deme circular stepping-stone models. Effect sizes were randomly sampled from a Gaussian distribution with variance 1; the value of (*t* − *t*_*W*_)*/t* was 0.1.

In contrast, the *Q*_*ST*_ distribution proposed by Koch (2019), in which *Q*_*ST*_ values are computed from simulated trait values drawn from a multivariate normal with covariance determined by mean coalescence times within and between demes, was well calibrated for polygenic traits regardless of number of demes and conservative for monogenic or oligogenic traits. Indeed, Supplementary Figures S10, S11, S12, and S13 show that Koch’s procedure performs well in all the settings we examined if 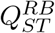 is used. As written, Koch’s procedure produces inflated type one error rates for 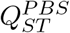 (Table S2). A modified version of Koch’s procedure would likely produce calibrated tests of 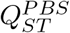, though we do not pursue this here. We caution that we used the true expected within- and between-deme coalescence times to calibrate Koch’s procedure, when in a realistic setting these times would need to be estimated.

Additionally, we tested a modification of the single-locus *F*_*ST*_ distribution strategy tested in Figure 3, in which we used the distribution of single-locus *F*_*ST*_ values, limiting only to common variants. Doing so typically produces well-calibrated type I error rates that are very similar to those produced by Koch’s method. Indeed, if allele-frequency changes among populations can be thought of as produced by drift well approximated by a multivariate normal distribution (Cavalli-Sforza et al., 1964; Nicholson et al., 2002; Berg and Coop, 2014), then we would expect single-locus *F*_*ST*_ to have the same distribution Koch proposed for *Q*_*ST*_ . (See supplementary text for more details on this claim.) For rare variants, allele-frequency change due to drift is not well approximated by a normal distribution— one reason is that because allele frequencies cannot drift below zero, the distribution of possible allele frequencies after drift is asymmetric. However, for sufficiently common variants and sufficiently short drift times, single-locus *F*_*ST*_ values might be expected to have a distribution similar to Koch’s proposal for neutral *Q*_*ST*_ . Supplementary Figures S6, S7, S8, and S9 show that the distribution of *F*_*ST*_ values for common alleles typically performs well as a null distribution for *Q*_*ST*_, so long as 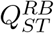 values are compared with 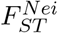 and 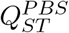 values are compared with 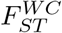.

For a summary of our findings in error rates in *Q*_*ST*_ –*F*_*ST*_ comparisons, see Figure 6. Supplementary Figure S14, S15, and Table S2 show type I error rate results in each demographic model with (*t* − *t*_*W*_)*/t* = 0.1, and Figure S16 shows results across different effect size distribution families.

**Figure 6.**
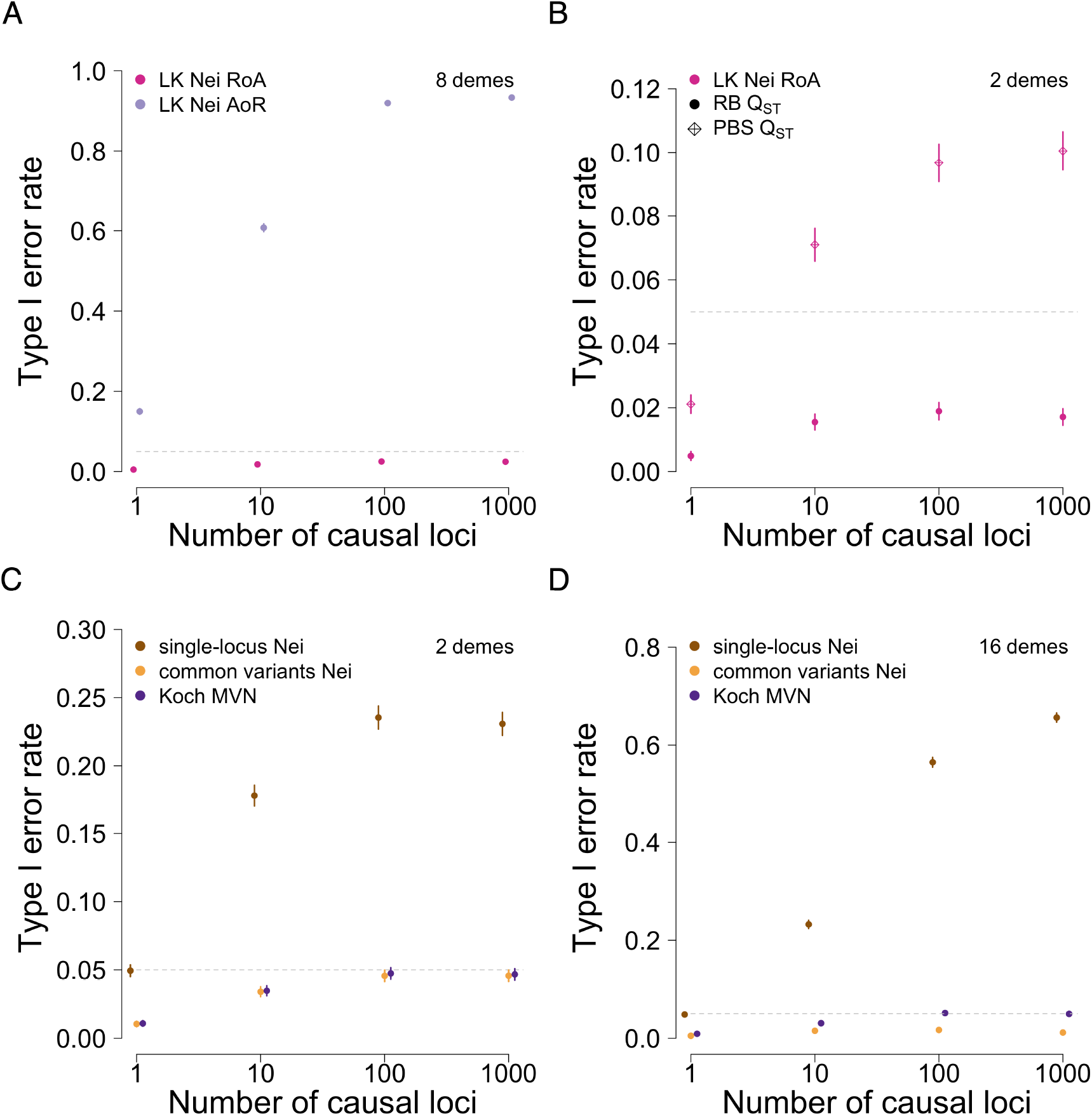
Summary of main results in terms of type I error rates. **A)** Ratio-of-averages estimates of genome-wide *F*_*ST*_ tend to produce calibrated or conservative type I error rates. In contrast, average-of-ratios *F*_*ST*_ is biased downward, causing elevated type I error rates when used to parameterize the Lewontin–Krakauer distribution. **B)** The versions of *F*_*ST*_ and *Q*_*ST*_ used should match in terms of their coalescent interpretations. Using 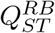 with 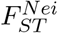 tends to produce calbrated or conservative results, as does using 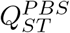 with 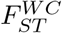. **C-D)** Using the full distribution of single-locus *F*_*ST*_ values produces calibrated tests for randomly chosen single-locus traits while anticonservative for polygenic traits. Using the distribution of single-locus *F*_*ST*_ values for common variants produces conservative *p* values. Koch’s (2019) procedure also produces calibrated *p* values for polygenic traits when 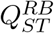 is used.

## 4 Discussion

We examined the effect of various choices for computing *Q*_*ST*_ and forming a null distribution on type I error rates in *Q*_*ST*_ –*F*_*ST*_ comparisons to detect local adaptation. In general, our results are all well explained if *Q*_*ST*_ and *F*_*ST*_ are viewed in terms of coalescent theory. That is, *Q*_*ST*_ –*F*_*ST*_ comparisons are well calibrated as tests of local adaptation if *Q*_*ST*_ is compared with a null distribution that approximates the distribution of the version of *Q*_*ST*_ chosen under a neutral coalescent process.

Although the distribution of *Q*_*ST*_ is sometimes argued not to depend on the number of loci that influence the trait, our simulations show that this is not quite true. Rather, the distribution of *Q*_*ST*_ differs for traits influenced by very small numbers of loci, generally being lower variance, and tends reach a limit as the number of loci becomes large. This behavior has been noticed previously (Edge and Rosenberg, 2015; Koch, 2019). In our simulations, polygenic traits lead to a higher-variance *Q*_*ST*_ distribution than monogenic or oligogenic traits, so using a *Q*_*ST*_ distribution calibrated for polygenic traits as a null will be conservative in tests of local adaptation. If a given trait is known to be monogenic, then one might argue that using the distribution of single-locus *F*_*ST*_ values is more appropriate, as suggested by Figure 3. However, in practice, we believe such a choice would often be inappropriate. Most monogenic traits that catch researchers’ interest for a *Q*_*ST*_ vs. *F*_*ST*_ test are likely to do so because they display substantial genetic variance, either within or between demes. Such ascertainment of traits on the basis of their variance makes them unlike rare variants, which will be the plurality of mutations observed in a sequencing study. Thus, if a trait is known to be monogenic, it might be more appropriate to conduct a test of local adaptation that conditions on its overall frequency.

We also find that whatever the method used, null distributions built from 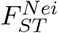 tend to work better when paired with 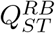, and null distributions built from 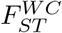 work best when paired with 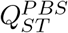, particularly when the number of demes is small. One way to understand this result is that neither 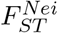 or 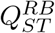 use Bessel’s correction when computing the among-population variance, whereas both 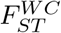 and 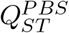 do use Bessel’s correction. Weaver (2016) also showed that both 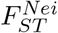 and 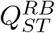 correspond to (*t* − *t*_*W*_)*/t*, where *t* is the average pairwise coalescence time for alleles drawn from the population at large, and *t*_*W*_ is the average pairwise coalescence time for alleles drawn at random from the same subpopulation. Similarly, 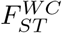 and 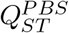correspond to (*t*_*B*_ − *t*_*W*_)*/t*_*B*_, where *t*_*B*_ is the average pairwise coalescence time for alleles drawn from different subpopulations. When 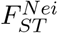 is used to develop a null distribution for 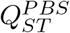, tests for local adaptation can be anti-conservative when the number of demes is small. This issue is subtle when the number of demes is large, but it is also easy to miss—indeed, in Koch’s (2019) paper, which presents the approach that performs best overall here, the distribution developed is most appropriate for 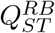, but it is compared with 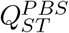 in simulations.

We find that in many settings, the Lewontin–Krakauer distribution provides an acceptable null distribution for *Q*_*ST*_ on polygenic traits, with calibrated or somewhat conservative type I error rates. However, it is important that the Lewontin–Krakauer distribution is parameterized by the correct version of *F*_*ST*_ . Specifically, in our simulations, the Lewontin– Krakauer distribution works best when parameterized by 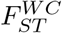 if 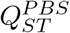 is the test statistic, and by 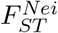 if 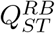 is the test statistic. Further, the genome-wide *F*_*ST*_ should be estimated via a ratio-of-averages approach—average-of-ratios estimators are biased downward, particularly if relatively rare variants are included, leading to excess type I errors in tests for local adaptation.

The one scenario we tested in which the Lewontin–Krakauer distribution consistently failed, even when appropriately parameterized, was in circular stepping-stone models with large numbers of demes. Spatial structure has previously been observed to lead to difficulties with the Lewontin–Krakauer distribution as a null distribution for *Q*_*ST*_ with large numbers of demes (Koch, 2019). However, in these scenarios, and in all others, we observed that Koch’s (2019) procedure produced calibrated type I error rates for polygenic traits when used as a null distribution for 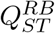. Though we did not pursue it explicitly, we also suspect that a slight modification of Koch’s procedure would produce calibrated type I error rates for 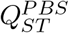 with small numbers of demes. Koch’s procedure computes *Q*_*ST*_ values by simulating genetic values for traits that obey a multivariate normal distribution with expectation zero and covariance determined by the average within- and between-deme coalescence times. Koch (2019) showed that this distribution is a good approximation for sufficiently polygenic traits with effect-size distributions that are not too heavy tailed. Here, we used the known coalescence time distributions to parameterize Koch’s procedure. However, this does not distinguish it much from other procedures we tested, as we simulated large numbers of neutral loci and thus generated very precise *F*_*ST*_ estimates.

Finally, we also tested use of the distribution of single-locus *F*_*ST*_ values as a null distribution for *Q*_*ST*_ . If all loci were used, this procedure produced calibrated type I errors for random monogenic traits (but see above), and badly anti-conservative tests for polygenic traits. However, limiting the single-locus *F*_*ST*_ values to those at loci with common minor alleles rescued the procedure for polygenic traits, causing it to perform well in every scenario tested. Our favored explanation for this is that drift at sufficiently common variants over short timescales can be approximated by a normal distribution (Nicholson et al., 2002; Berg and Coop, 2014). Thus, for common variants, the distribution of allele frequencies among subpopulations might be well approximated by the multivariate normal distribution developed by Koch (2019). Presumably the procedure for defining “common” variants for inclusion should depend to some degree on the type of population structure observed, but we do not pursue this question here.

Our work here focused specifically on the “evolutionary” variation in neutral *Q*_*ST*_ . That is, we assumed that we had access to the genetic values of the trait (also called breeding values) for a large number of individuals per deme, as well as genotypes at a large number of selectively neutral loci for each individual. Thus, we focused on variation caused by the evolutionary-genetic process and did not consider the effect of uncertainty in estimating the within- and among-deme genetic variance in the trait, and in estimating *F*_*ST*_ . In real applications, these other considerations are important (Whitlock, 2008), but it is also important to consider the “evolutionary” variation in its own right, as we have done here, because it exists regardless of study design or precision of measurement.

In recent years, alternatives to *Q*_*ST*_ –*F*_*ST*_ comparisons have been developed that take advantage of more information about population structure than provided by *F*_*ST*_ alone (Ovaskainen et al., 2011; Berg and Coop, 2014; Josephs et al., 2019). Koch’s (2019) method for developing a null distribution for *Q*_*ST*_ can be seen as part of this family of extensions, as it uses the set of mean within- and between-deme coalescence times to produce a null distribution for *Q*_*ST*_ rather using the value of *F*_*ST*_ itself. Such methods can produce more powerful or better calibrated tests of local adaptation in some cases. However, the properties of *Q*_*ST*_ –*F*_*ST*_ comparisons that we study here are still important. One reason is that common-garden studies, which are necessary for rigorous interpretation (Brommer, 2011; Schraiber and Edge, 2024), are difficult and time-consuming to perform, and many have been performed at substantial effort and expense, not all of which will have retained the data necessary to perform a reanalysis with a more modern method. There is thus value in ensuring that the lessons learned from common-garden studies are robust. To do so, it would be fruitful to consider the types of markers used in many common-garden *Q*_*ST*_ –*F*_*ST*_ comparisons—in many cases, data from microsatellites or RADseq—from the coalescent perspective used here. For example, estimates of *F*_*ST*_ from microsatellites are often lower than for other markers (Jakobsson et al., 2013), which might be expected to lead to *Q*_*ST*_ values that spuriously indicate local adaptation (Edelaar et al., 2011). Measures of genetic differentiation at microsatellites designed to estimate the same function of coalescence times as Nei’s *F*_*ST*_ —for example, Slatkin’s *R*_*ST*_ (Slatkin, 1995)—might provide a way forward in such cases if their assumptions are met. As such, the coalescent perspective on neutral quantitative-trait differentiation (Whitlock, 1999; Koch, 2019) can inform both new analyses and reanalyses of valuable archival data on local adaptation.

## 5 Acknowledgments

We thank members of the Edge, Mooney, and Pennell labs for comments that improved this work. Funding was provided by NIH grant R35GM137758 to MDE.

## 6 Code Accessibility

Code used to run and analyze the simulations in this study is available at https://github.com/junjianliu/qst_fst.

