## Supplementary text and figures for "Error rates in *Q_ST_ –F_ST_* comparisons depend on genetic architecture and estimation procedures"

### Supplementary text for "Error rates in $Q_{ST}$–$F_{ST}$ comparisons depend on genetic architecture and estimation procedures"

Junjian J. Liu and Michael D. Edge

Compiled on October 29, 2024

In the supplement, we provide mathematical details to support the results in the main text.

#### S1 Values of $t$, $t_W$, and $t_B$ in various demographic models

##### S1.1 Setting $(t - t_W)/t$ and identifying demographic parameters

Slatkin (1991) showed that $F_{ST}^{Nei}$ has a low-mutation-rate limit of

$$\frac{t - t_W}{t}, \tag{S1}$$

where $t$ is mean coalescence time of two alleles chosen uniformly at random from the total population, and $t_W$ is the mean coalescence time of two random alleles within the same subpopulation. In this section, we abuse notation and refer to this function of coalescence times as $F_{ST}^{Nei}$. We sought to set $(t - t_W)/t$ at various desired values under each of several demographic models: a star-like split mode, a "balanced" split model, a "caterpillar" split model, and island model, and a circular stepping-stone model. Here, we provide expressions for $t$ and $F_{ST}^{Nei}$ under each of these models. In all our models, we assume constant population sizes that are equal in all subpopulations, as well as equal-sized samples from each subpopulation. (We set $N_e = 1000$ in all demes in practice.)

In models involving population trees, we assume that all branches have the same constant population size $N_e$. Thus, in all split models, $t_W = 2N_e$. For a star-like population tree, there is a single split time $t_s$ at which time all populations split from an ancestral population. If the number of individuals sampled per subpopulation is large, then two random lineages are drawn from the same subpopulation with probability $1/d$ and from different subpopulations with probability $(d-1)/d$. (This is only approximate—the probability two random lineages are drawn from the same population is $(2n-1)/(2nd-1)$, where $n$ is the number of diploid individuals sampled per deme, due to sampling without replacement. In our simulations, $n$ is large enough that $(2n-1)/(2nd-1) \approx 1/d$, and we use the approximation.) If the two lineages are drawn from different populations, their time to coalesce is $2N_e + t_s$ generations. Thus, using the law of total expectation, the average time for a random pair of lineages to coalesce in the starlike model is

$$t_{(starlike)} = 2N_e + \frac{(d-1)t_s}{d}.$$

$F_{ST}^{Nei}$ is therefore

$$F_{ST(starlike)}^{Nei} = \frac{1}{1 + \frac{2N_e d}{(d-1)t_s}}.$$

In the balanced model, we assume that all splits are $t_s$ generations apart; i.e. the most recent splits are $t_s$ generations before the present, the next most recent are $2t_s$ generations before the present, etc. This assumption leads to

$$t_{(balanced)} = 2N_e + \frac{(d \log_2 d - d + 1)t_s}{d}$$

and therefore

$$F_{ST(balanced)}^{Nei} = \frac{1}{1 + \frac{2N_e d}{(d \log_2 d - d + 1)t_s}}.$$

The "balanced" tree requires that the number of demes is a power of 2.

In a the caterpillar model, we again assume that subsequent splits on the population tree are $t_s$ generations apart, leading to

$$t_{(caterpillar)} = 2N_e + \frac{(d-1)(2d-1)t_s}{3d}$$

and

$$F_{ST(caterpillar)}^{Nei} = \frac{1}{1 + \frac{6N_e d}{(d-1)(2d-1)t_s}}.$$

Similarly, for a $d$-deme migration model with a local effective population size $N_e$ and a total migration rate $m$ evenly distributed from a deme to its receivers, $t_W$ is the same in every deme and equal to

$$t_{W(migration)} = 2N_e d,$$

while the values for $t$ in different demographic models are

$$t_{(island)} = 2N_e d + \frac{(d-1)^2}{2md},$$

$$t_{(circular)} = 2N_e d + \frac{(d+2)(2d-1) - 3d}{24m}.$$

Plugging in $t$ and $t_W$ values for each model into the expression of $F_{ST}^{Nei}$ (eq. S1) gives

$$F_{ST(island)}^{Nei} = \frac{1}{1 + \frac{4N_e md^2}{(d-1)^2}},$$

$$F_{ST(circular)}^{Nei} = \frac{1}{1 + \frac{24N_e md}{d^2-1}}.$$

Here, we assumed that the number of demes in the circular stepping stone model was even.

For all models, we picked values of either $t_s$ or $m$ to achieve $(t - t_W)/t$ values of 1/50, 1/10, and 1/4.

#### S1.2 Calculating $(t_B - t_W)/t_W$

Slatkin (1993) showed that $F_{ST}^{WC}$ has a low-mutation-rate limit of

$$F_{ST}^{WC} = \frac{t_B - t_W}{t_B}, \tag{S2}$$

where $t_B$ is the mean coalescence time of two random alleles from two different subpopulations. Slatkin also gave the relationship between the pairwise mean coalescence times $t$, $t_B$, and $t_W$. Equation 10 in Slatkin (1995) gives

$$t = \frac{2n(d-1)}{2nd-1} t_B + \frac{2n-1}{2nd-1} t_W, \tag{S3}$$

where $n$ is the number of sampled diploid individuals in each subpopulation. However, if $n >> 1$, this is very closely approximated by

$$t = \frac{d-1}{d} t_B + \frac{1}{d} t_W, \tag{S4}$$

which is implicit in the previous subsection. We use this approximation to relate $(t - t_W)/t$ and $(t_B - t_W)/t_B$, since at the sample sizes we simulate, the approximation error is extremely small.

Under the approximation in equation S4, some algebra lets us write

$$\frac{t - t_W}{t} = \frac{(d-1)(t_B - t_W)}{(d-1)t_B + t_W}.$$

If we again abuse notation and define $F_{ST}^{Nei} = (t - t_W)/t$, then we obtain

$$\frac{t_B - t_W}{t_B} = \frac{F_{ST}^{Nei} d}{F_{ST}^{Nei} + d - 1},$$

which we can use to identify the estimand of $F_{ST}^{WC}$ (i.e. $(t_B - t_W)/t_B$) given the values of $(t - t_W)/t$ and the number of demes (see Table S1).

| $F_{ST}^{Nei}$ | $F_{ST}^{WC}$ | | | |
|---|---|---|---|---|
| | 2D | 4D | 8D | 16D |
| 1/50 | 2/51 | 4/151 | 8/351 | 16/751 |
| 1/10 | 2/11 | 4/31 | 8/71 | 16/151 |
| 1/4 | 2/5 | 4/13 | 8/29 | 16/61 |

Table S1: $F_{ST}$ estimands in models with various number of demes

### S2 Koch's multivariate normal distribution and allele-frequency differences among subpopulations

In this section, we sketch an informal argument that for loci with sufficiently common minor allele frequencies in the ancestral population, the distribution of single-locus $F_{ST}$ values approximately follows the distribution for $Q_{ST}$ on additive, polygenic traits identified by Koch (2019). Specifically, we consider a neutral model and assume allele-frequency drift can be modeled using a normal distribution (Cavalli-Sforza et al., 1964; Nicholson et al., 2002; Berg and Coop, 2014).

Berg & Coop (2014) proposed a model of neutral allele-frequency change at a biallelic locus in which

$$\vec{p} \sim \mathcal{MVN}(p_0\vec{1}, p_0(1-p_0)\mathbf{F}),$$

where $\vec{p}$ is a vector of allele frequencies in the $d$ subpopulations at the present, $p_0$ is the allele frequency in a shared ancestral population, and $\mathbf{F}$ is a matrix describing the covariance of allele-frequency drift across subpopulations. Under this model, Berg & Coop showed that the subpopulation-mean genetic values of a neutrally evolving polygenic trait are distributed as

$$\vec{G} \sim \mathcal{MVN}(\mu\vec{1}, 2V_A\mathbf{F}),$$

where $\vec{G}$ is a vector of subpopulation-mean genetic values, $\mu$ is the mean genetic value in the shared ancestral population, $V_A$ is the additive genetic variance, and $\mathbf{F}$ is the same matrix appearing in the distribution of $\vec{p}$. (Koch's procedure entails simulating from a multivariate normal in which $\mathbf{F}$ is parameterized by mean within- and between-subpopulation coalescence times.) Thus, under the multivariate-normal model of allele-frequency drift assumed by Berg & Coop, the covariance matrix for the subpopulation-mean genetic values for the trait is proportional to the covariance matrix of subpopulation allele frequencies.

If all subpopulation sample sizes are the same, $F_{ST}^{Nei}$ at a locus can be estimated as

$$F_{ST}^{Nei} = \frac{\frac{1}{d}\sum_i (p_i - \bar{p})^2}{\frac{1}{d}\sum_i (p_i - \bar{p})^2 + \frac{1}{d}\sum_i p_i(1-p_i)}.$$

Similarly, if all subpopulation samples sizes are the same,

$$Q_{ST}^{Koch} = \frac{\frac{1}{d}\sum_i (G_i - \bar{G})^2}{\frac{1}{d}\sum_i (G_i - \bar{G})^2 + \frac{1}{d}\sum_i V_{W(i)}},$$

where $V_{W(i)}$ is the within-subpopulation variance of the genetic values in subpopulation $i$. Noticing that the within-subpopulation variance in a draw of a single allelic indicator variable is $p_i(1 - p_i)$, we can see that these expressions are both estimated fractions of variance due to among-subpopulation differences in two variables of the same distribution family with proportional covariance matrices. Thus, they should have approximately the same distribution.

In practice, the multivariate normal model of allele-frequency drift is likely only a good fit for sufficiently common alleles that have been drifting for a sufficiently short amount of time (Paris et al., 2019). This may explain why we observe similar distributions for $Q_{ST}$ and single-locus $F_{ST}$ values only if we restrict to $F_{ST}$ for common variants.

| | single-locus Nei | single-locus WC | LK Nei RoA | LK Nei AoR | LK WC RoA | LK WC AoR | Koch MVN | common variants Nei | common variants WC |
|---|---|---|---|---|---|---|---|---|---|
| | $Q_{ST}^{RB}$ | | | | | | | | |
| 2D star-like | 0.2308 | 0.0957 | 0.0171 | 0.3105 | 0 | 0.1694 | 0.0467 | 0.0457 | 0.0039 |
| 4D star-like | 0.31 | 0.1879 | 0.0229 | 0.6265 | 0.0031 | 0.5168 | 0.0446 | 0.0186 | 0.0034 |
| 8D star-like | 0.4589 | 0.3605 | 0.0246 | 0.9333 | 0.0095 | 0.9113 | 0.0456 | 0.0183 | 0.0078 |
| 16D star-like | 0.6562 | 0.5933 | 0.0295 | 0.9991 | 0.0147 | 0.9989 | 0.0501 | 0.0115 | 0.0056 |
| 4D balanced | 0.3158 | 0.2058 | 0.0401 | 0.6101 | 0.011 | 0.5067 | 0.0495 | 0.039 | 0.0138 |
| 8D balanced | 0.4337 | 0.3602 | 0.0564 | 0.9044 | 0.0346 | 0.8772 | 0.05 | 0.0299 | 0.0187 |
| 16D balanced | 0.6387 | 0.5941 | 0.0796 | 0.9957 | 0.0616 | 0.9948 | 0.0506 | 0.0272 | 0.0206 |
| 4D caterpillar | 0.3268 | 0.2115 | 0.0323 | 0.6166 | 0.0075 | 0.5166 | 0.0481 | 0.0248 | 0.0074 |
| 8D caterpillar | 0.4358 | 0.3551 | 0.0389 | 0.9071 | 0.0198 | 0.8813 | 0.0447 | 0.0202 | 0.012 |
| 16D caterpillar | 0.6115 | 0.5564 | 0.0564 | 0.9967 | 0.0391 | 0.9962 | 0.0506 | 0.0234 | 0.0156 |
| 2D island | 0.1555 | 0.0454 | 0.0198 | 0.2277 | 0.0004 | 0.0949 | 0.0451 | 0.0531 | 0.0059 |
| 4D island | 0.1452 | 0.0653 | 0.0221 | 0.3583 | 0.0038 | 0.2344 | 0.05 | 0.0346 | 0.0103 |
| 8D island | 0.1166 | 0.068 | 0.0277 | 0.5 | 0.0104 | 0.411 | 0.0516 | 0.0277 | 0.0116 |
| 16D island | 0.0787 | 0.0527 | 0.0298 | 0.671 | 0.0184 | 0.6118 | 0.0516 | 0.0262 | 0.016 |
| 4D circular | 0.1331 | 0.0641 | 0.0288 | 0.3426 | 0.0055 | 0.2199 | 0.0472 | 0.0431 | 0.0135 |
| 8D circular | 0.1429 | 0.0986 | 0.057 | 0.469 | 0.0315 | 0.4006 | 0.0487 | 0.0424 | 0.0239 |
| 16D circular | 0.1596 | 0.1356 | 0.1038 | 0.6116 | 0.0856 | 0.5769 | 0.0495 | 0.0485 | 0.0379 |
| | $Q_{ST}^{PBS}$ | | | | | | | | |
| 2D star-like | 0.396 | 0.2308 | 0.1005 | 0.4763 | 0.0017 | 0.3239 | 0.1605 | 0.1588 | 0.0457 |
| 4D star-like | 0.446 | 0.3107 | 0.0646 | 0.7252 | 0.0177 | 0.6356 | 0.11 | 0.0546 | 0.0187 |
| 8D star-like | 0.5499 | 0.4602 | 0.0556 | 0.9523 | 0.0227 | 0.9364 | 0.0881 | 0.042 | 0.0183 |
| 16D star-like | 0.7144 | 0.6598 | 0.0446 | 0.9993 | 0.0286 | 0.9991 | 0.0744 | 0.0248 | 0.0119 |
| 4D balanced | 0.4334 | 0.3168 | 0.0831 | 0.706 | 0.0323 | 0.6202 | 0.0996 | 0.0811 | 0.039 |
| 8D balanced | 0.5121 | 0.4349 | 0.0848 | 0.9313 | 0.0533 | 0.9083 | 0.076 | 0.0459 | 0.0301 |
| 16D balanced | 0.6824 | 0.6411 | 0.1004 | 0.9972 | 0.0788 | 0.9965 | 0.0639 | 0.0364 | 0.0272 |
| 4D caterpillar | 0.4474 | 0.3275 | 0.079 | 0.7131 | 0.0251 | 0.6269 | 0.1089 | 0.0641 | 0.0248 |
| 8D caterpillar | 0.5211 | 0.4368 | 0.0692 | 0.9336 | 0.0369 | 0.9121 | 0.0773 | 0.0371 | 0.0203 |
| 16D caterpillar | 0.6691 | 0.6141 | 0.0776 | 0.9977 | 0.0556 | 0.9971 | 0.069 | 0.0318 | 0.0236 |
| 2D island | 0.3102 | 0.1555 | 0.0983 | 0.3912 | 0.0019 | 0.2345 | 0.1546 | 0.1708 | 0.0531 |
| 4D island | 0.2588 | 0.1453 | 0.0675 | 0.4891 | 0.0173 | 0.3652 | 0.1177 | 0.0883 | 0.0347 |
| 8D island | 0.1803 | 0.1171 | 0.0546 | 0.5922 | 0.025 | 0.5067 | 0.092 | 0.0544 | 0.0277 |
| 16D island | 0.1089 | 0.0793 | 0.0461 | 0.7297 | 0.0289 | 0.6782 | 0.0787 | 0.0403 | 0.0265 |
| 4D circular | 0.2341 | 0.1336 | 0.0723 | 0.4732 | 0.0221 | 0.348 | 0.1059 | 0.0973 | 0.0432 |
| 8D circular | 0.1924 | 0.1434 | 0.0862 | 0.5415 | 0.0538 | 0.4735 | 0.0756 | 0.07 | 0.0426 |
| 16D circular | 0.1869 | 0.1605 | 0.1259 | 0.6476 | 0.1031 | 0.6161 | 0.0618 | 0.0602 | 0.0486 |

Table S2: Type I error rates from 10,000 simulated traits in $Q_{ST}$–$F_{ST}$ comparisons for traits with 1000 causal loci. $(t - t_W)/t = 0.1$ WC = Weir & Cockerham; LK = Lewontin–Krakauer; RoA = Ratio of Averages; AoR = Average of Ratios.

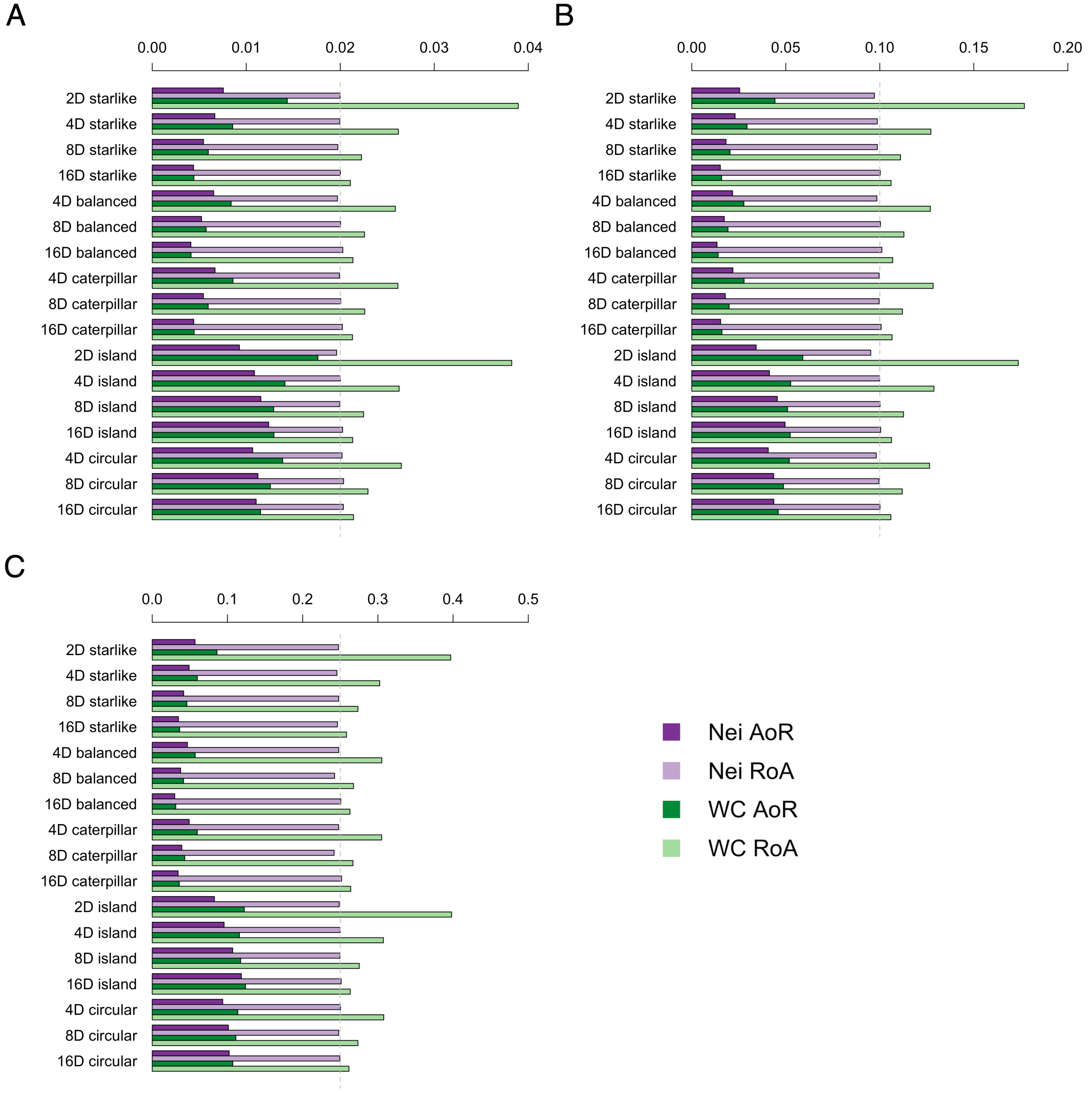


Figure S1: $F_{ST}$ **estimates under various demographic models.** We simulated independent coalescent trees and used the branch lengths to approximate three joint allele frequency spectra per demographic model. We set $(t - t_W)/t$ (corresponding to $F_{ST}^{Nei}$) to (A) 0.02, (B) 0.1, or (C) 0.25. This figure shows ratio-of-average and average-of ratios estimates of genome-wide $F_{ST}^{Nei}$ and $F_{ST}^{WC}$ for all variants.

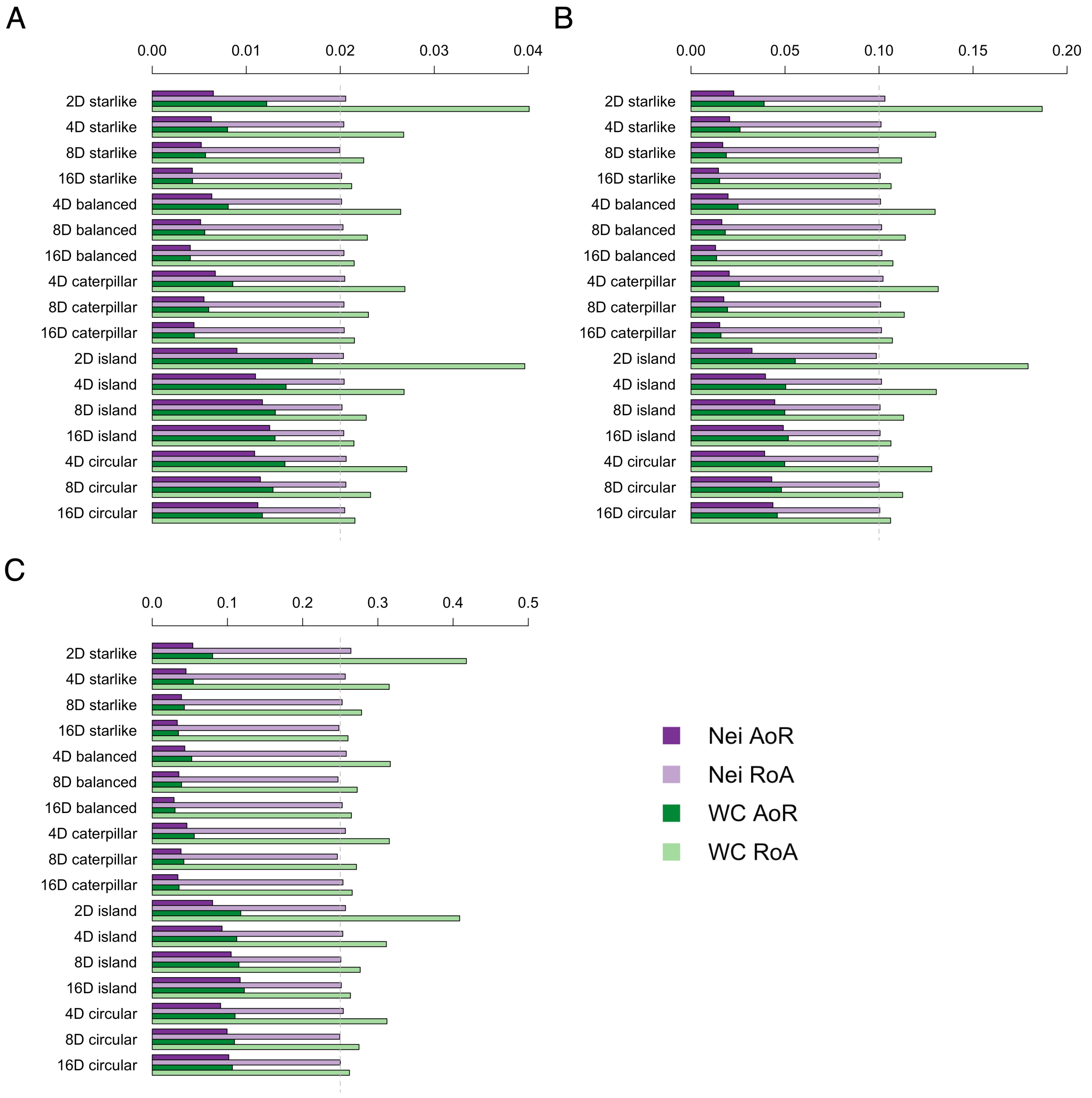


Figure S2: $F_{ST}$ **estimates under various demographic models using loci with common minor alleles in a specific subpopulation** We simulated independent coalescent trees and used the branch lengths to approximate three joint allele frequency spectra per demographic model. We set $(t - t_W)/t$ (corresponding to $F_{ST}^{Nei}$) to (A) 0.02, (B) 0.1, or (C) 0.25. This figure shows ratio-of-average and average-of ratios estimates of genome-wide $F_{ST}^{Nei}$ and $F_{ST}^{WC}$ for variants ascertained to be common (with a frequency greater than 0.05) in a specific subpopulation.

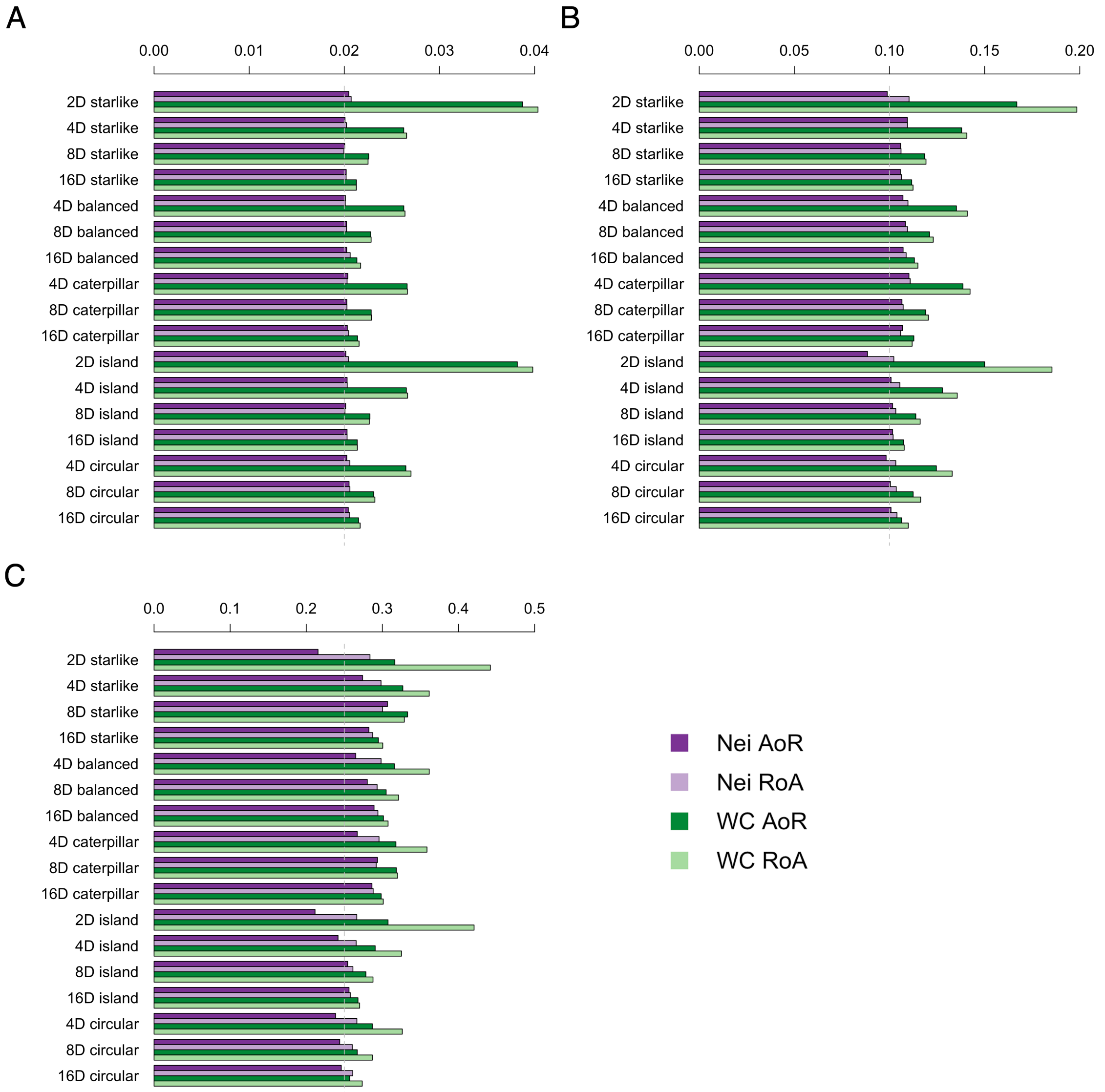


Figure S3: $F_{ST}$ **estimates under various demographic models using loci with common minor alleles in the combined population.** We simulated independent coalescent trees and used the branch lengths to approximate three joint allele frequency spectra per demographic model. We set $(t - t_W)/t$ (the estimand of $F_{ST}^{Nei}$) to (A) 0.02, (B) 0.1, or (C) 0.25. This figure shows ratio-of-average and average-of ratios estimates of genome-wide $F_{ST}^{Nei}$ and $F_{ST}^{WC}$ for variants ascertained to be common (with a frequency greater than 0.05) across subpopulations.

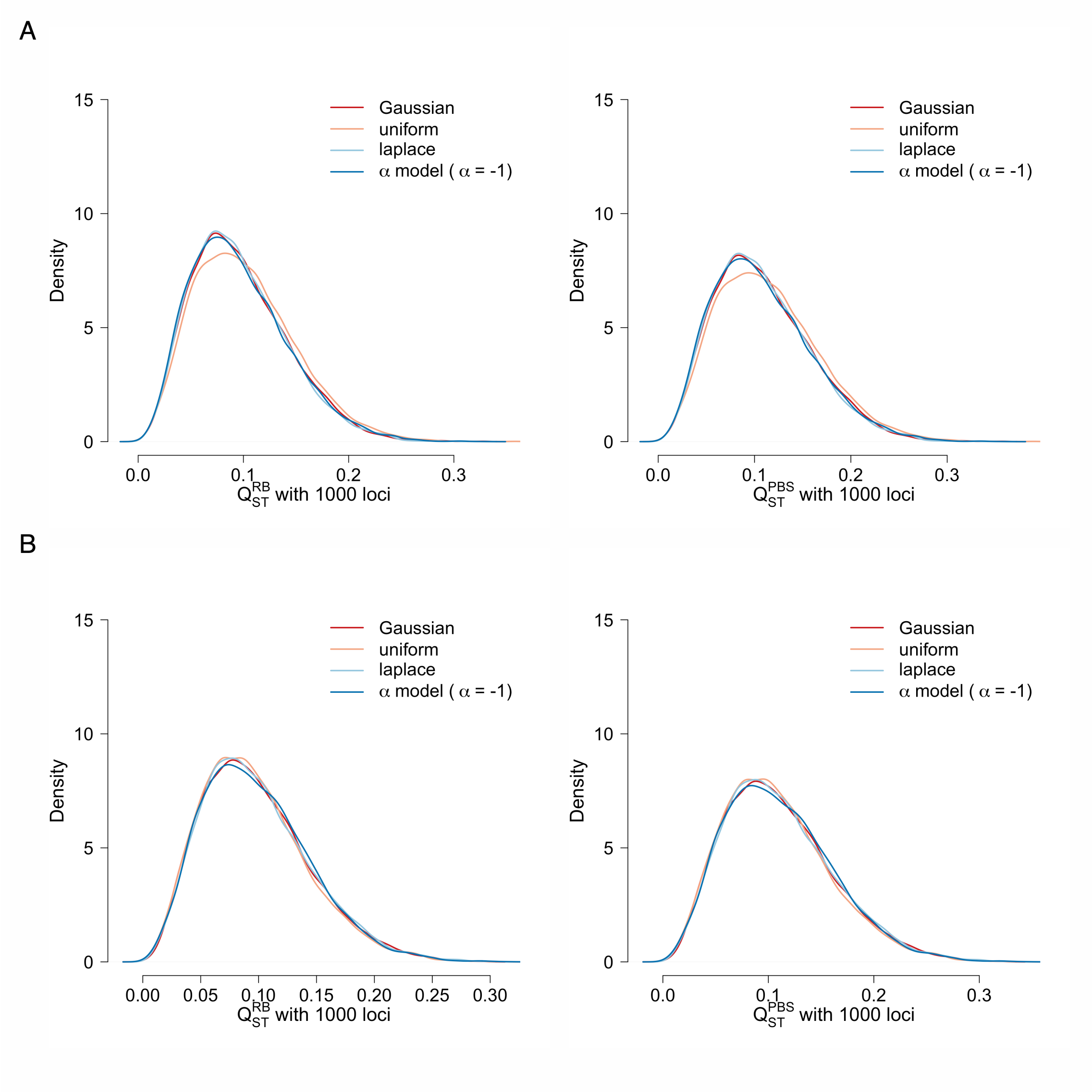


Figure S4: $Q_{ST}$ **distributions for 1000 causal loci with effect sizes from different distribution families.** Effect sizes were drawn from Gaussian, uniform, and Laplace distributions with expectation 0 and variance 1. We also tested effect sizes drawn from an "alpha model" with $\alpha = -1$ (an allele-frequency-dependent Gaussian distribution in which the effect-size standard deviation is inversely proportional to $\sqrt{\bar{p}(1-\bar{p})}$, where $\bar{p}$ is the mean allele frequency across the total population). The panel shows results in (A) an eight-deme star-like split model and (B) an eight-deme island model.

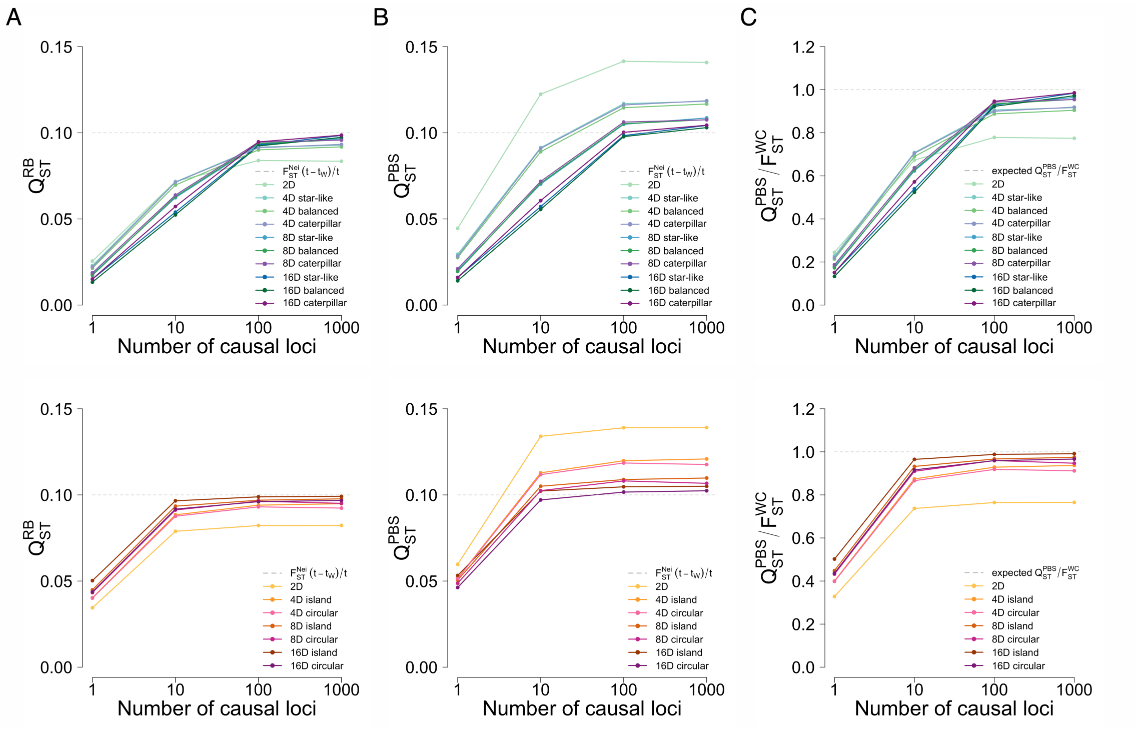


Figure S5: **The behavior of mean $Q_{ST}$ estimates in all demographic models.** Effect sizes were randomly sampled from a Gaussian distribution with variance 1 to generate genetic values. Mean $Q_{ST}$ estimates were calculated across 1000 simulated traits with $(t - t_W)/t$ (i.e. the function of coalescent times estimated by $F_{ST}^{Nei}$) equal to 0.1. The curves in each panel show the behavior of (A) $Q_{ST}^{RB}$ in all split and migration models, (B) $Q_{ST}^{PBS}$ in all split and migration models, (C) $Q_{ST}^{PBS}/F_{ST}^{WC}$ in all split and migration models.

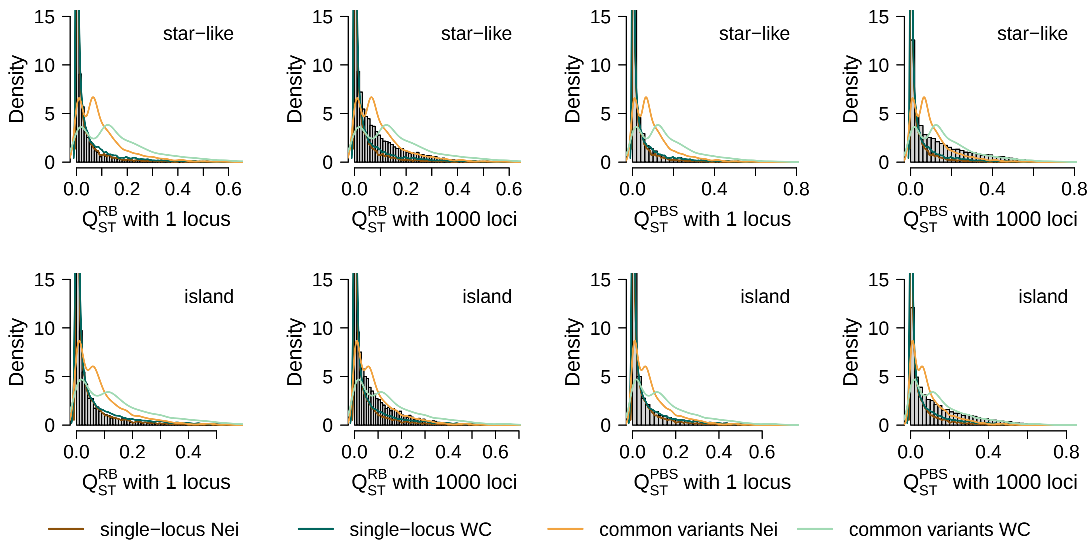


Figure S6: **Single-locus $F_{ST}$ density curves vs. $Q_{ST}$ distributions across genetic architectures: two-deme star-like split and island models.** We compared four null distributions (the single-locus $F_{ST}^{Nei}$ and $F_{ST}^{WC}$ density curves using all variable loci, and curves using common variants whose allele frequency is greater than 0.05 in the combined population only) with the neutral $Q_{ST}^{RB}$ and $Q_{ST}^{PBS}$ distributions. Each $Q_{ST}$ distribution included 10,000 traits with 1 or 1000 causal loci. The panels show the results for a two-deme star-like split model and a two-deme island model. Effect sizes were randomly sampled from a Gaussian distribution. $(t - t_W)/t = 0.1$.

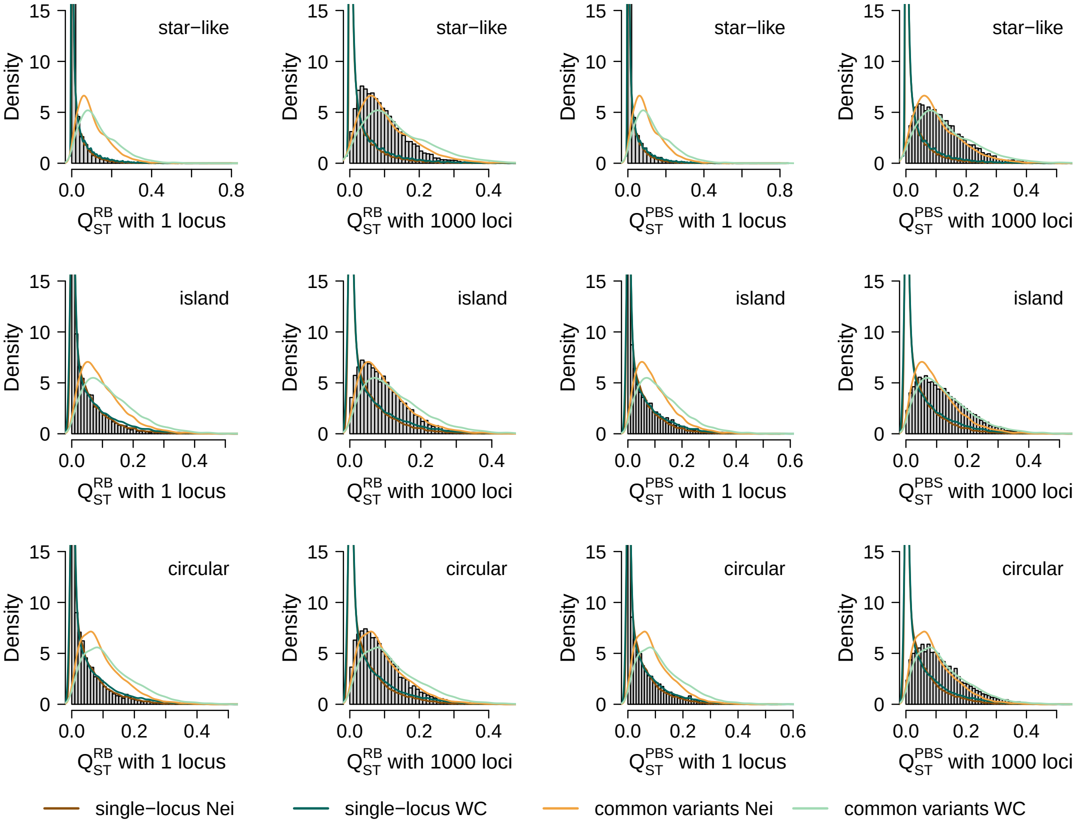


Figure S7: **Single-locus $F_{ST}$ density curves vs. $Q_{ST}$ distributions across genetic architectures: four-deme star-like split, island, and circular stepping-stone models.** We compared four null distributions (the single-locus $F_{ST}^{Nei}$ and $F_{ST}^{WC}$ density curves using all variable loci, and curves using common variants whose allele frequency is greater than 0.05 only) with the neutral $Q_{ST}^{RB}$ and $Q_{ST}^{PBS}$ distributions. Each $Q_{ST}$ distribution included 10,000 traits with 1 or 1000 causal loci. The panels show the results for a four-deme star-like split model, a four-deme island model, and a four-deme circular stepping-stone model. Effect sizes were randomly sampled from a Gaussian distribution. $(t - t_W)/t = 0.1$.

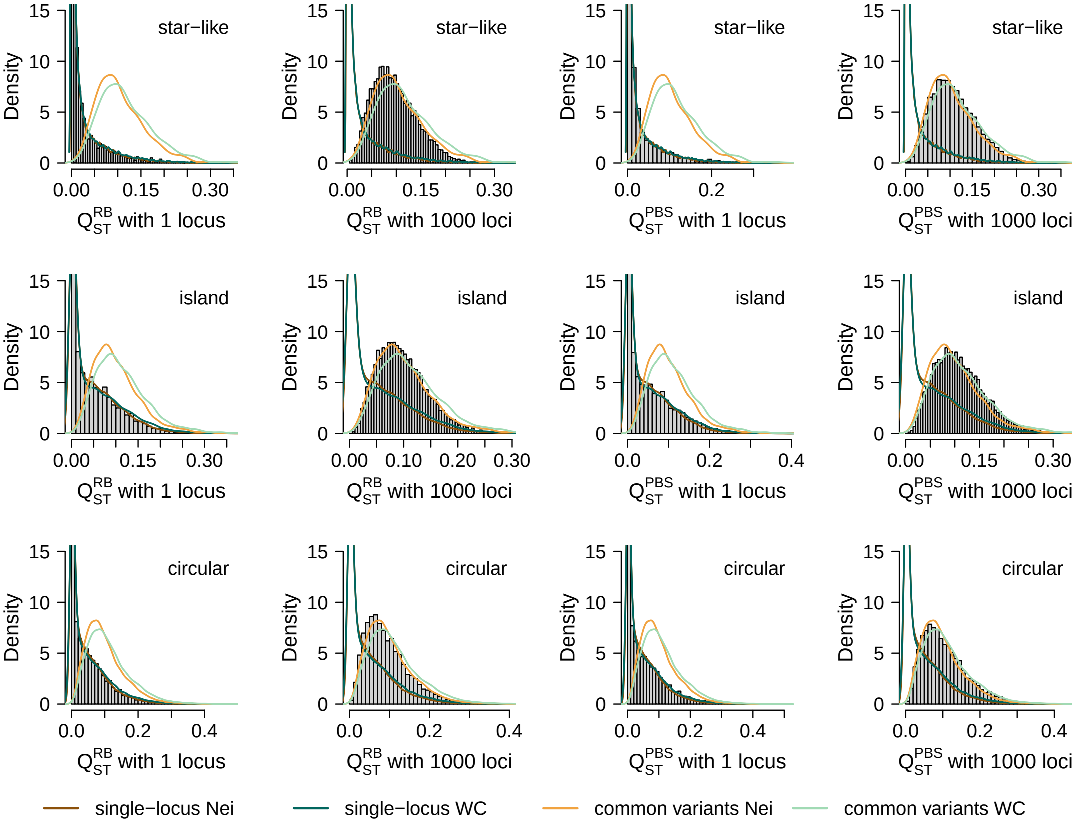


Figure S8: **Single-locus $F_{ST}$ density curves vs. $Q_{ST}$ distributions across genetic architectures: eight-deme star-like split, island, and circular stepping-stone models.** We compared four null distributions (the single-locus $F_{ST}^{Nei}$ and $F_{ST}^{WC}$ density curves using all variable loci, and curves using common variants whose allele frequency is greater than 0.05 only) with the neutral $Q_{ST}^{RB}$ and $Q_{ST}^{PBS}$ distributions. Each $Q_{ST}$ distribution included 10,000 traits with 1 or 1000 causal loci. The panels show the results for an eight-deme star-like split model, an eight-deme island model, and an eight-deme circular stepping stone model. Effect sizes were randomly sampled from a Gaussian distribution. $(t - t_W)/t = 0.1$.

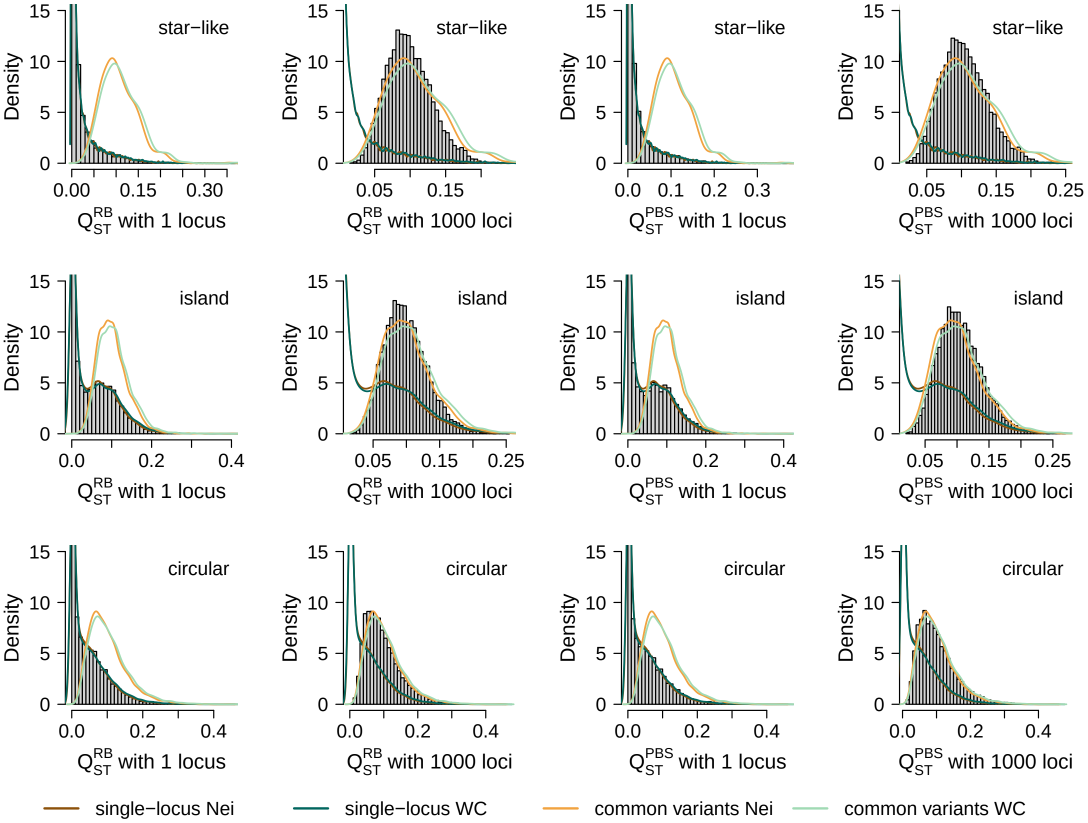


Figure S9: **Single-locus $F_{ST}$ density curves vs. $Q_{ST}$ distributions across genetic architectures: sixteen-deme star-like split, island, and circular stepping-stone models.** We compared four null distributions (the single-locus $F_{ST}^{Nei}$ and $F_{ST}^{WC}$ density curves using all variable loci, and curves using common variants whose allele frequency is greater than 0.05 only) with the neutral $Q_{ST}^{RB}$ and $Q_{ST}^{PBS}$ distributions. Each $Q_{ST}$ distribution included 10,000 traits with 1 or 1000 causal loci. The panels show the results for a sixteen-deme star-like split model, a sixteen-deme island model, and a sixteen-deme circular stepping-stone model. Effect sizes were randomly sampled from a Gaussian distribution. $(t - t_W)/t = 0.1$.

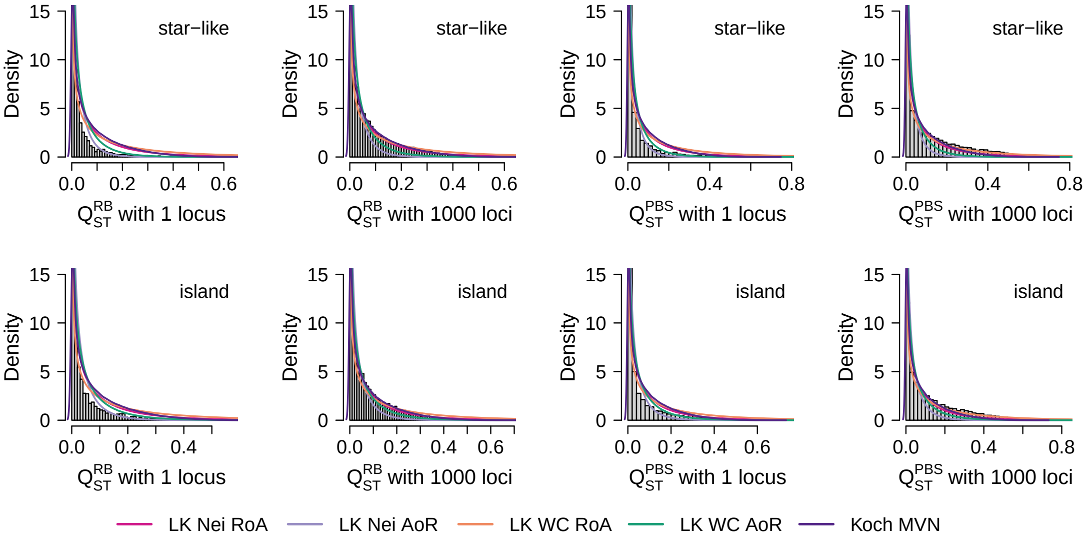


Figure S10: **Lewontin–Krakauer null and Koch's null vs. $Q_{ST}$ distributions across genetic architectures: two-deme star-like split and island models.** We compared five null distributions (the Lewontin–Krakauer distribution parameterized by either ratio-of-average or average-of ratios estimates of genome-wide $F_{ST}^{Nei}$ or $F_{ST}^{WC}$, and Koch's multivariate normal distribution) with the neutral $Q_{ST}^{RB}$ and $Q_{ST}^{PBS}$ distributions. Each $Q_{ST}$ distribution included 10,000 traits with 1 or 1000 causal loci. The panels show the results for a two-deme star-like split model and a two-deme island model. Effect sizes were randomly sampled from a Gaussian distribution. $(t - t_W)/t = 0.1$.

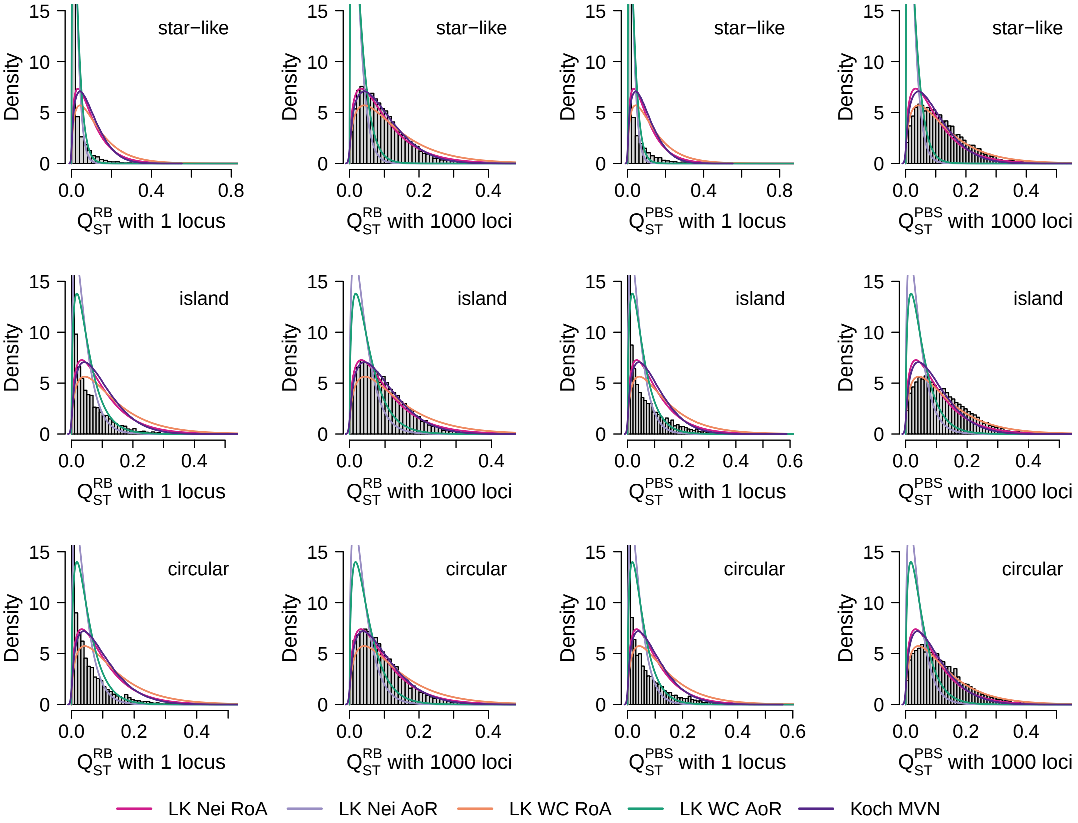


Figure S11: **Lewontin–Krakauer null and Koch's null vs. $Q_{ST}$ distributions across genetic architectures: four-deme star-like split, island, and circular stepping-stone models.** We compared five null distributions (the Lewontin–Krakauer distribution parameterized by either ratio-of-average or average-of ratios estimates of genome-wide $F_{ST}^{Nei}$ or $F_{ST}^{WC}$, and Koch's multivariate normal distribution) with the neutral $Q_{ST}^{RB}$ and $Q_{ST}^{PBS}$ distributions. Each $Q_{ST}$ distribution included 10,000 traits with 1 or 1000 causal loci. The panels show the results for a four-deme star-like split model, a four-deme island model, and a four-deme circular stepping-stone model. Effect sizes were randomly sampled from a Gaussian distribution. $(t - t_W)/t = 0.1$.

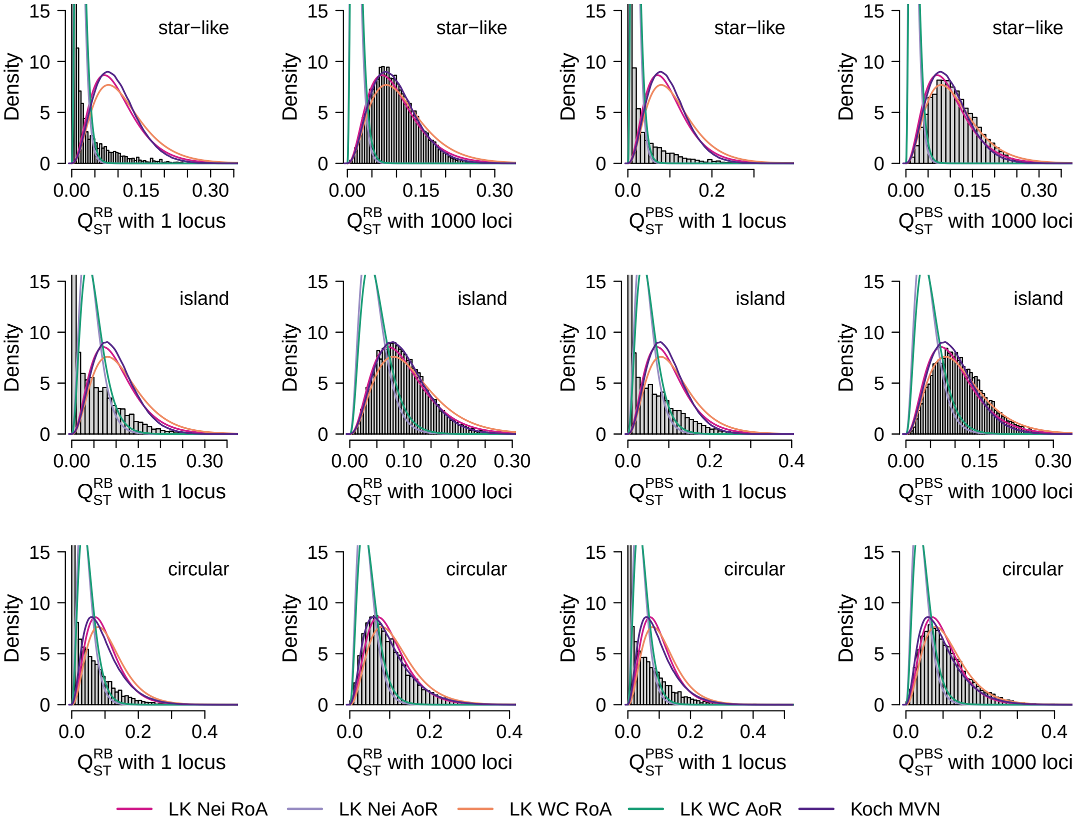


Figure S12: **Lewontin–Krakauer null and Koch's null vs. $Q_{ST}$ distributions across genetic architectures: eight-deme star-like split, island, and circular stepping-stone models.** We compared five null distributions (the Lewontin–Krakauer distribution parameterized by either ratio-of-average or average-of ratios estimates of genome-wide $F_{ST}^{Nei}$ or $F_{ST}^{WC}$, and Koch's multivariate normal distribution) with the neutral $Q_{ST}^{RB}$ and $Q_{ST}^{PBS}$ distributions. Each $Q_{ST}$ distribution included 10,000 traits with 1 or 1000 causal loci. The panels show the results for an eight-deme star-like split model, an eight-deme island model, and an eight-deme circular stepping stone model. Effect sizes were randomly sampled from a Gaussian distribution. $(t - t_W)/t = 0.1$.

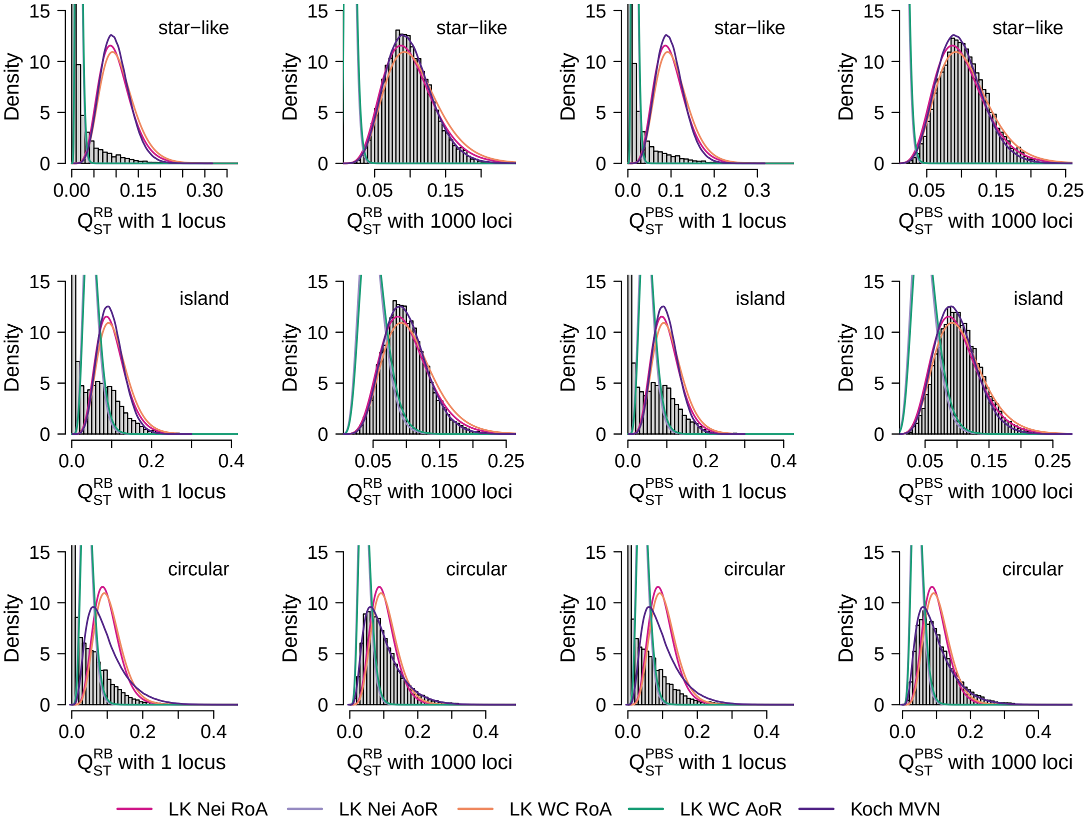


Figure S13: **Lewontin–Krakauer null and Koch's null vs. $Q_{ST}$ distributions across genetic architectures: sixteen-deme star-like split, island, and circular stepping-stone models.** We compared five null distributions (the Lewontin–Krakauer distribution parameterized by either ratio-of-average or average-of ratios estimates of genome-wide $F_{ST}^{Nei}$ or $F_{ST}^{WC}$, and Koch's multivariate normal distribution) with the neutral $Q_{ST}^{RB}$ and $Q_{ST}^{PBS}$ distributions. Each $Q_{ST}$ distribution included 10,000 traits with 1 or 1000 causal loci. The panels show the results for a sixteen-deme star-like split model, a sixteen-deme island model, and a sixteen-deme circular stepping-stone model. Effect sizes were randomly sampled from a Gaussian distribution. $(t - t_W)/t = 0.1$.

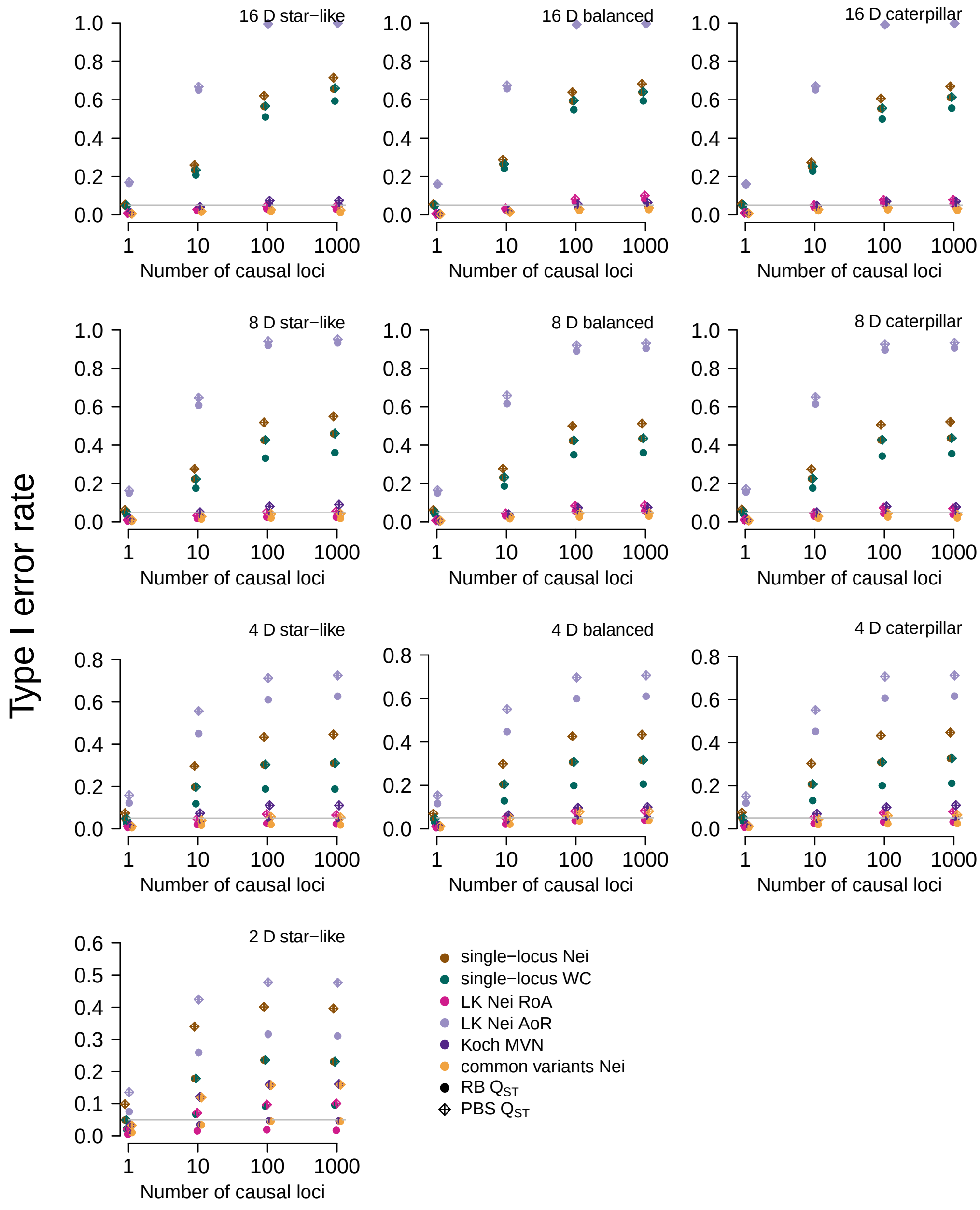


Figure S14: **Selected type I error rates in $Q_{ST}$–$F_{ST}$ comparisons of star-like, balanced, and caterpillar split models.** Effect sizes were randomly sampled from a Gaussian distribution with variance 1; $(t - t_W)/t = 0.1$.

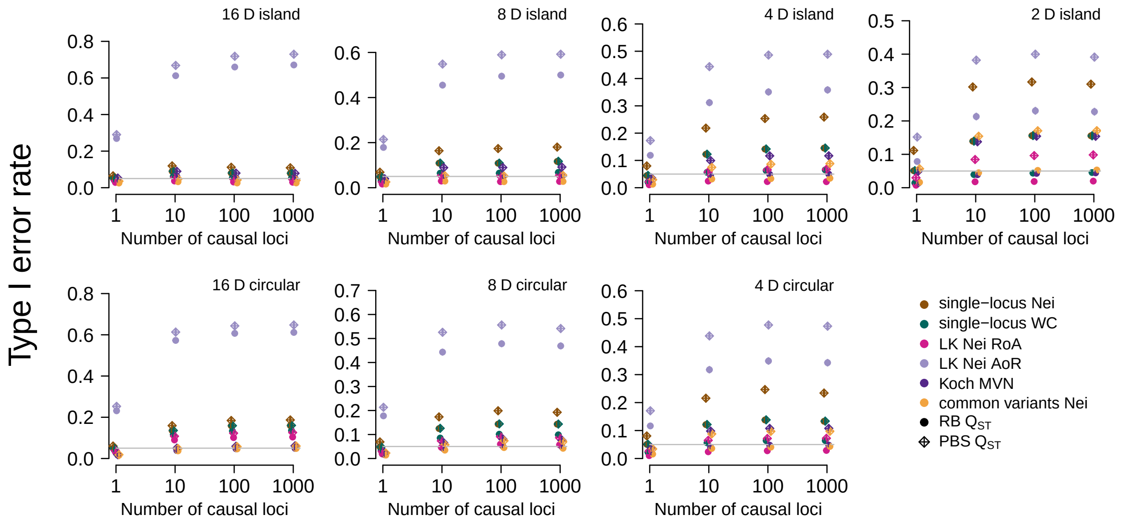


Figure S15: **Selected type I error rates in $Q_{ST}$–$F_{ST}$ comparisons of island and circular stepping-stone models.** Effect sizes were randomly sampled from a Gaussian distribution with variance 1; $(t - t_W)/t = 0.1$.

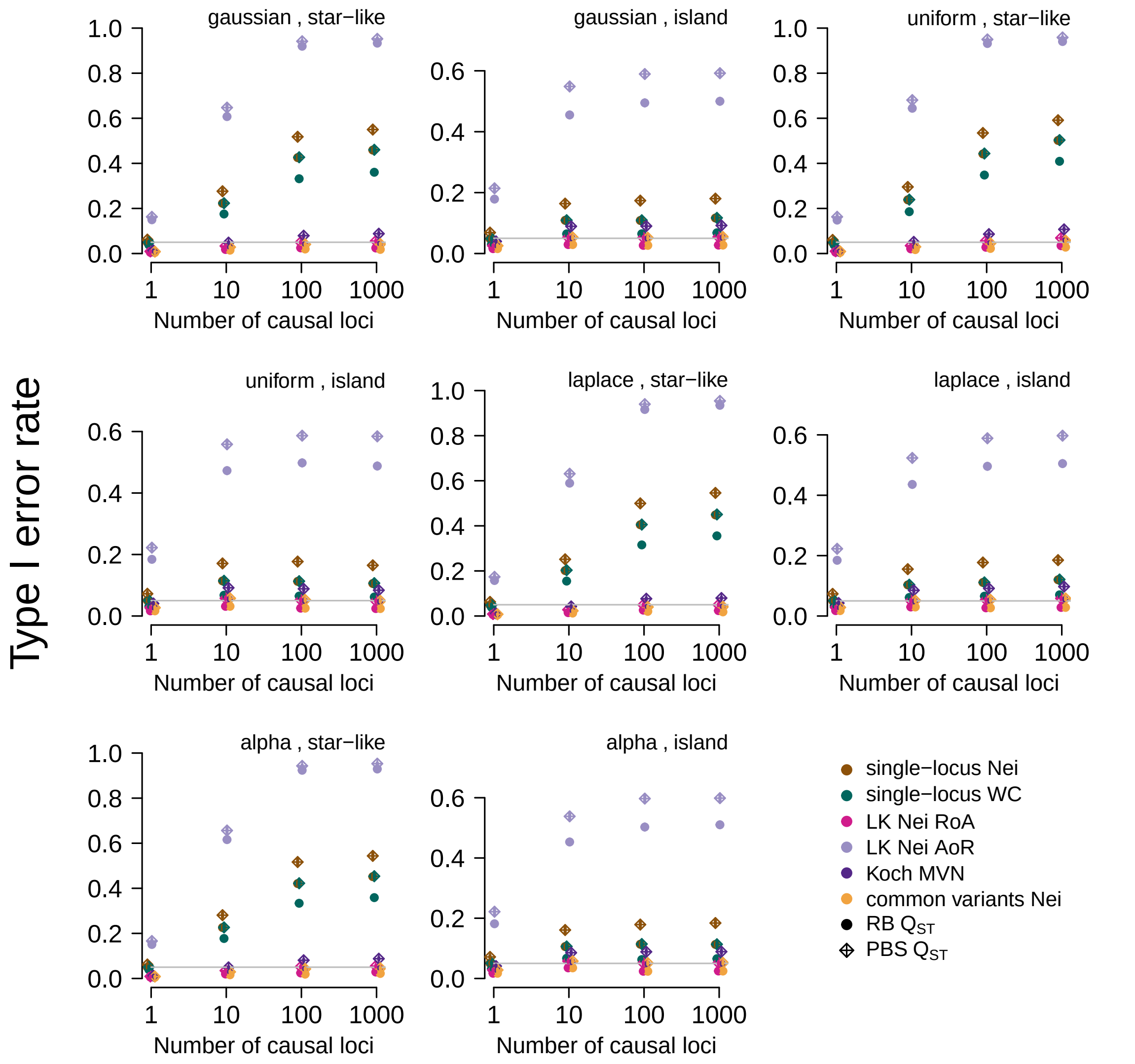


Figure S16: **Selected type I error rates in $Q_{ST}$–$F_{ST}$ comparisons across effect size distribution families: eight-deme star-like split and island models.** Effect sizes were drawn from Gaussian, uniform, and laplace distributions with expectation 0 and variance 1. We also tested effect sizes drawn from an "alpha model" with $\alpha = -1$ (an allele-frequency-dependent Gaussian distribution in which the effect-size standard deviation is inversely proportional to $\sqrt{\bar{p}(1-\bar{p})}$, where $\bar{p}$ is the mean allele frequency across the total population). $(t - t_W)/t = 0.1$.
